# Leaf starch metabolism sets the phase of stomatal rhythm

**DOI:** 10.1101/2022.10.07.511256

**Authors:** Adrianus J. Westgeest, Myriam Dauzat, Thierry Simonneau, Florent Pantin

## Abstract

In leaves of C_3_ and C_4_ plants, stomata open during the day to favour CO_2_ entry for photosynthesis, and close at night to prevent inefficient transpiration of water vapour. The circadian clock paces rhythmic stomatal movements throughout the diel (24-h) cycle. Leaf transitory starch is also thought to regulate the diel stomatal movements, yet the underlying mechanisms across time (key moments) and space (relevant leaf tissues) remains elusive. Here, we developed PhenoLeaks, a pipeline to analyse the diel dynamics of transpiration, and used it to screen a series of Arabidopsis mutants impaired in starch metabolism. We detected a sinusoidal, endogenous rhythm of transpiration that overarches days and nights. We uncovered that a number of severe mutations in starch metabolism affect the endogenous rhythm through a phase shift, resulting in delayed stomatal movements throughout the daytime and reduced stomatal preopening during the night. Nevertheless, analysis of tissue-specific mutations revealed that neither guard-cell nor mesophyll-cell starch metabolism are strictly required for normal diel patterns of transpiration. We propose that leaf starch influences the timing of transpiration rhythm through an interplay between the clock and sugars across tissues, while the energetic effect of starch-derived sugars is usually non-limiting for endogenous stomatal movements.

**One-sentence summary:** The PhenoLeaks pipeline for monitoring diel transpiration dynamics reveals that leaf starch metabolism sets the timing of the endogenous stomatal rhythm.

## INTRODUCTION

Leaf stomata are tiny pores surrounded by pairs of guard cells that control carbon gain and water loss in plants (Hetherington and Woodward, 2003). In C_3_ and C_4_ plants, stomata open every day to promote atmospheric CO_2_ uptake when light is available to power photosynthesis, but this also generates transpiration of water vapour from the inner leaf tissues into the atmosphere. When the light dims, stomata close to prevent inefficient water loss (Vialet-Chabrand et al., 2017). At night, stomata are essentially closed, yet progressive preopening is usually observed before dawn (Caird et al., 2007). Overall, the ability of stomata to track fast changes in irradiance or maintain low transpiration throughout the night enhances plant water use efficiency (Caird et al., 2007; Lawson and Blatt, 2014; Coupel-Ledru et al., 2016; Vialet-Chabrand et al., 2017; Papanatsiou et al., 2019; Eyland et al., 2021). As water becomes increasingly scarce with climate change, tailoring stomatal dynamics during day and night is a promising strategy to breed crops for drought tolerance. Stomatal movements ultimately result from osmotically-driven changes in turgor pressure of the guard cells, with feedbacks from the epidermal pavement cells and the inner leaf tissues (Lawson and Matthews, 2020; Blatt et al., 2022; Nieves-Cordones et al., 2022). Although the instantaneous effects of light fluctuations on the guard cells are relatively well understood, we lack a mechanistic framework depicting the dynamics of stomatal movements across the diel (24-h) cycle in the context of a whole leaf that is subjected to variations of both external and internal cues.

While the diel pattern of stomatal movement is primarily driven by light availability, it also has an endogenous rhythmic component that is controlled by the circadian clock (Hubbard and Webb, 2015). In day-night cycles alternating constant light and darkness, stomata start to open several hours before the end of the night, continue to open until the middle of the day, and then engage to close during the afternoon. This endogenous rhythm, *i.e.* that is observed in binary day/night cycles without other change in the environment, is under circadian control. For instance, in Arabidopsis, a circadian stomatal rhythm under continuous light is observed in wild-type plants, but is disturbed or even abolished in mutants defective in the circadian clock (Somers et al., 1998; Sothern et al., 2002; Dodd et al., 2004, 2005; Hassidim et al., 2017; Dakhiya and Green, 2019). The circadian rhythmicity of stomata connects with the physiology of the plant. It is more robust in a whole leaf than in an isolated epidermis (Gorton et al., 1989) and is the main driver of the circadian rhythm in photosynthesis (Hennessey and Field, 1991; Gorton et al., 1993). Moreover, despite a strong circadian control, the endogenous stomatal rhythm also shows high flexibility to the environment. During the day, in Arabidopsis plants placed in constant light, the endogenous stomatal rhythm vary in timing and amplitude depending on whether plants have been entrained in constant or fluctuating light (Matthews et al., 2018). In wheat, weak blue light enhances the amplitude of the daytime endogenous rhythm and this effect is respiration-dependent (Vialet-Chabrand et al., 2021). Decades of research have also attempted to decrypt stomatal dynamics in darkness, since the first observation one century ago that stomata preopen throughout the night (Loftfield, 1921). The effect of the photoperiod has been studied extensively (Schwabe, 1952; Mansfield and Heath, 1961, 1963, 1964; Martin and Meidner, 1971, 1972, 1975; Kana and Miller, 1977), eventually hinting at a role for a photosynthetic product (Holmes and Klein, 1986). More recently in *Vicia faba*, it was shown that nighttime preopening is abolished when plants are shaded on the preceding day, and starts earlier when they are given elevated CO_2_ (Easlon and Richards, 2009), suggesting a connection with carbon metabolism. Thus, metabolic cues may counteract, potentiate or interact with the clock to regulate stomatal dynamics across the whole diel cycle.

Starch metabolism and associated sugars could be involved in the diel control of stomatal movements, yet this hypothesis has remained underexplored. While it was early noticed that stomata of the starch-free mutant *pgm* do not preopen at night (Lascève et al., 1997), recent research rather focused on the role of starch and sugar metabolism in light-induced stomatal opening (Flütsch and Santelia, 2021). In leaf chloroplasts, a fraction of the sugars produced in the day by photosynthesis is accumulated as transitory starch, which is broken down in the following night (Stitt and Zeeman, 2012; Streb and Zeeman, 2012). This diel process of daytime storage and nighttime breakdown, which buffers the diel variations of metabolizable sugars against the fluctuations in light availability, occurs mainly in the mesophyll cells. In the guard cells, the turnover of starch is slightly out of phase, with starch synthesis persisting in the early night, and starch degradation accelerating in the early day (Santelia and Lunn, 2017). Rapid guard-cell starch degradation during the first hour of the day is controlled by non-photosynthetic levels of blue light and generates glucose to support stomatal opening (Horrer et al., 2016; Flütsch et al., 2020b). When photosynthesis rises with light, the guard cells import most of their sugars (glucose and sucrose) from the mesophyll; such import further enhances stomatal opening and replenishes the guard-cell starch pool simultaneously (Flütsch et al., 2020b, 2020a). Yet, the general role of sugars in stomatal movements is still debated (Daloso et al., 2016a; Santelia and Lawson, 2016; Daloso et al., 2017; Granot and Kelly, 2019; Lawson and Matthews, 2020; Flütsch and Santelia, 2021). Sugars may trigger stomatal opening by providing the signal, energy or osmotica required for turgor build-up of the guard cells (Outlaw and Manchester, 1979; Poffenroth et al., 1992; Daloso et al., 2015, 2016b; Flütsch et al., 2020a, 2020b). Sugars may also evoke stomatal closure either in the apoplast as osmotica that pull water out of the guard cells (Lu et al., 1997), or in the guard cells through hexokinase-mediated sugar sensing (Kelly et al., 2013, 2019). It has been proposed that sugars induce different effects depending on the carbon status, which covaries with the time of the day, so that sugars promote opening under starvation, like at dawn, but trigger closure upon satiation, like in the afternoon (Flütsch and Santelia, 2021). This working model has remained difficult to test, yet, because the available data for sugar concentration lack spatiotemporal resolution, and because uncoupling variation in carbon status from time of the day is challenging. While many mutants of the starch metabolism are available with different impacts on sugar content at different times of the diel cycle and in different cell types, we reasoned that obtaining their diel transpiration patterns would provide insight into the spatiotemporal hierarchy of events that connect carbon metabolism and guard cell dynamics throughout the diel cycle.

Here, we developed a phenotyping pipeline to screen the diel transpiration dynamics of a series of Arabidopsis mutants differentially impaired in the starch metabolic pathway (Figure 1). Our rationale was i) to phenotype a collection of starch mutants over the diel cycle so as to identify the most prominent patterns, ii) to challenge these dynamics by subjecting the plants to environments that modify carbon status, and iii) to develop a quantitative framework allowing us to translate the transpiration dynamics into relevant biological parameters. We found that severe mutations in starch synthesis (*pgm*, *isa1*) or breakdown (*sex1*, *mex1*) affect the endogenous rhythm of transpiration through a phase shift, which results in delayed stomatal movements throughout the daytime and/or reduced stomatal preopening at night. However, affecting starch synthesis in the mesophyll but not in the guard cells (*pgi*), blocking starch degradation in the guard cells only (*amy3 bam1*), or impairing starch degradation in both tissues (*bam1 bam3*) did not substantially affect the transpiration dynamics, suggesting that minimal starch turnover in either tissue is sufficient for the guard cells to phase properly. We concluded that leaf starch metabolism influences the endogenous dynamics of transpiration indirectly by modulating the sugar dynamics across tissues, most likely through an interaction with the circadian clock – while an energetic effect of starch-derived sugars only emerges upon severe starvation. Our work sheds light on the regulation of transpiration dynamics, providing new tracks to breed crops for more efficient use of water.

**Figure 1.**
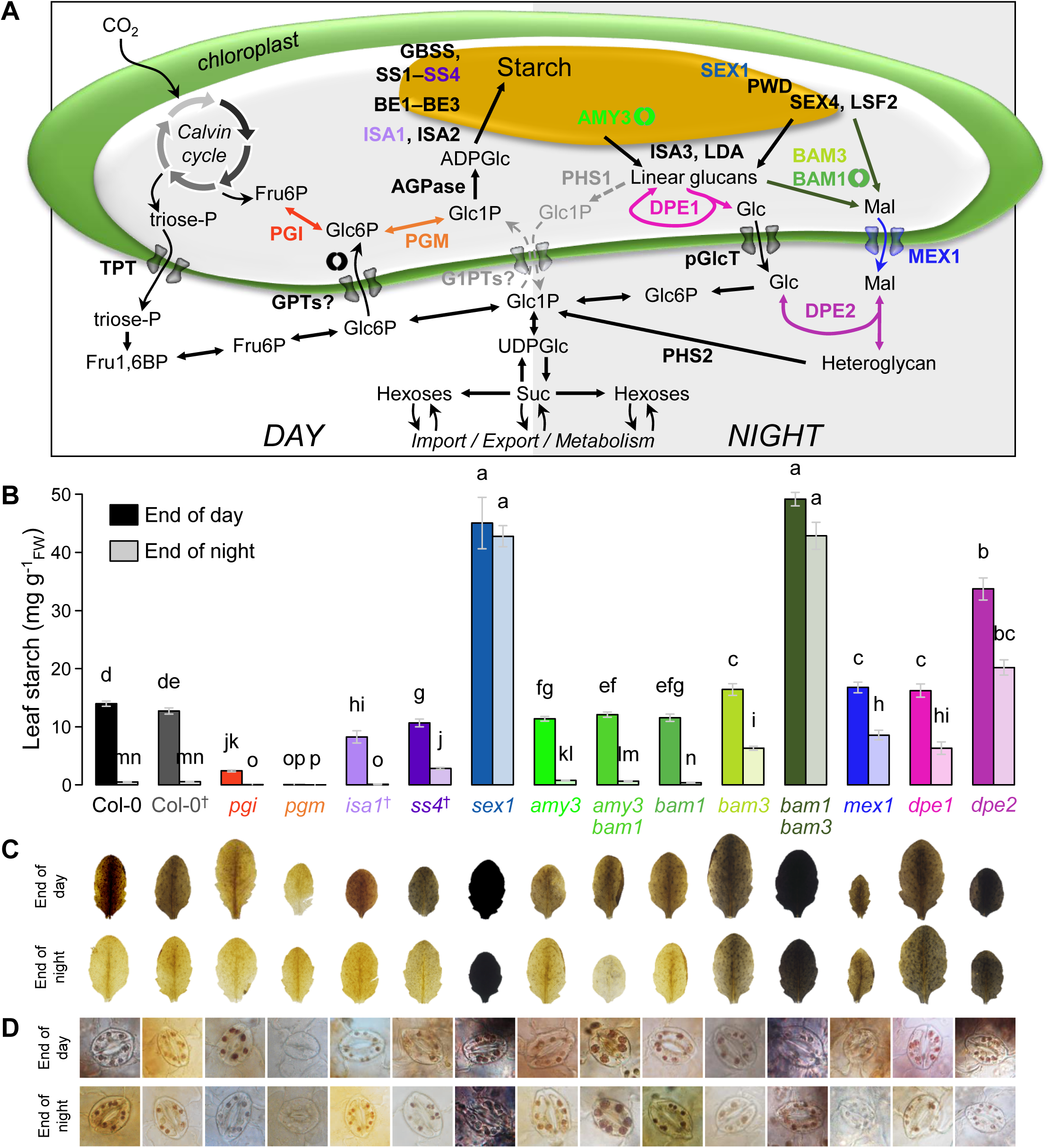
Characterization of the starch patterns in the starch mutants. **(A)** Schematic view of the pathway of transitory starch in Arabidopsis leaves (adapted from Stitt and Zeeman, 2012; Santelia and Lunn, 2017). In mesophyll cells, starch (yellow shape) is synthetized in the chloroplast (green shape) during the day (left) and broken down at night (right), while it is slightly out of phase in the guard cells. Steps that are specific to the guard cells are depicted by a stomata symbol, but the function of many of the other proteins in the guard cells has not been yet documented. Note that BAM3 is specific to the mesophyll, and that BAM1 can compensate for its absence. Steps that are likely to be minor are indicated in grey, dashed arrows. Uncertain chloroplastic transporters are indicated with a question mark. **(B)** Leaf starch content determined at the end of the day and night periods in control conditions. Error bars are means ± SE. Letters denote significant differences after a Kruskal-Wallis test (α = 5%) followed by multiple comparisons of ranks. **(C)** and **(D)** Iodine staining of starch on entire leaves **(C)** or individual guard cells **(D)** at the end of the day (top rows) or night (bottom rows). The dagger in Col-0^†^, *ss4*^†^ and *isa1*^†^ indicates that these lines harbour a luciferase reporter that was not used in this study (Methods).

## RESULTS

### Developing PhenoLeaks for high-throughput analysis of the diel transpiration dynamics

Monitoring transpiration throughout the diel cycle is critically informative on the dynamics of the processes underlying water loss, but remains a phenotyping bottleneck. We developed PhenoLeaks, a phenotyping pipeline to analyse transpiration dynamics from a temporal series of gravimetrical data. We first adapted a high-throughput phenotyping platform fitted with a mobile scale (Granier et al., 2006) and developed novel scripts so as to monitor the diel kinetics of transpiration on 150 Arabidopsis plants during several days with a 30-min resolution in controlled environmental conditions (Figure 2A; Methods). We first present transpiration of the wild type Col-0 in four independent experiments at the same daytime irradiance (180 µmol m^-2^ s^-1^) and atmospheric CO_2_ (400 ppm), yet with slightly different conditions such as the vapour pressure deficit (VPD, which varied between 0.75 and 1 kPa but was constant days and nights within experiments), the plant age (between 29 and 41 days after sowing), and the Phenopsis automaton used for growing or assaying the plants (Supplemental Table S1). Across all experiments, we observed temporal patterns that could be ascribed to rapid stomatal movements at the day/night transitions and to slower, endogenous stomatal rhythms otherwise (Figure 2B). Stomata opened rapidly when the light switched on, continued to open until the middle of the day and then partly closed until the end of the day. When the light switched off, stomata closed rapidly and, after about 1-3 h, started to reopen gradually throughout the night. Nonetheless, stomatal preopening was barely observed in experiment #1 (which was the only assay where temperature was cooler in the nighttime than in the daytime). Other variations were noticeable between experiments, such as the baseline, the diel amplitude or the timing when daytime transpiration reaches its maximum (Figure 2B). Unsurprisingly, differences in the transpiration baseline were linked to the VPD, which physically drives transpiration. However, differences in the dynamics (amplitude and timing) could not be consistently attributed to an evident source of variation (Supplemental Table S1, Supplemental Figure S1 for statistical comparisons of parameters described thereafter). This stressed the importance of monitoring different plants simultaneously when comparing *e*.*g*. different genotypes, which was done thereafter.

**Figure 2.**
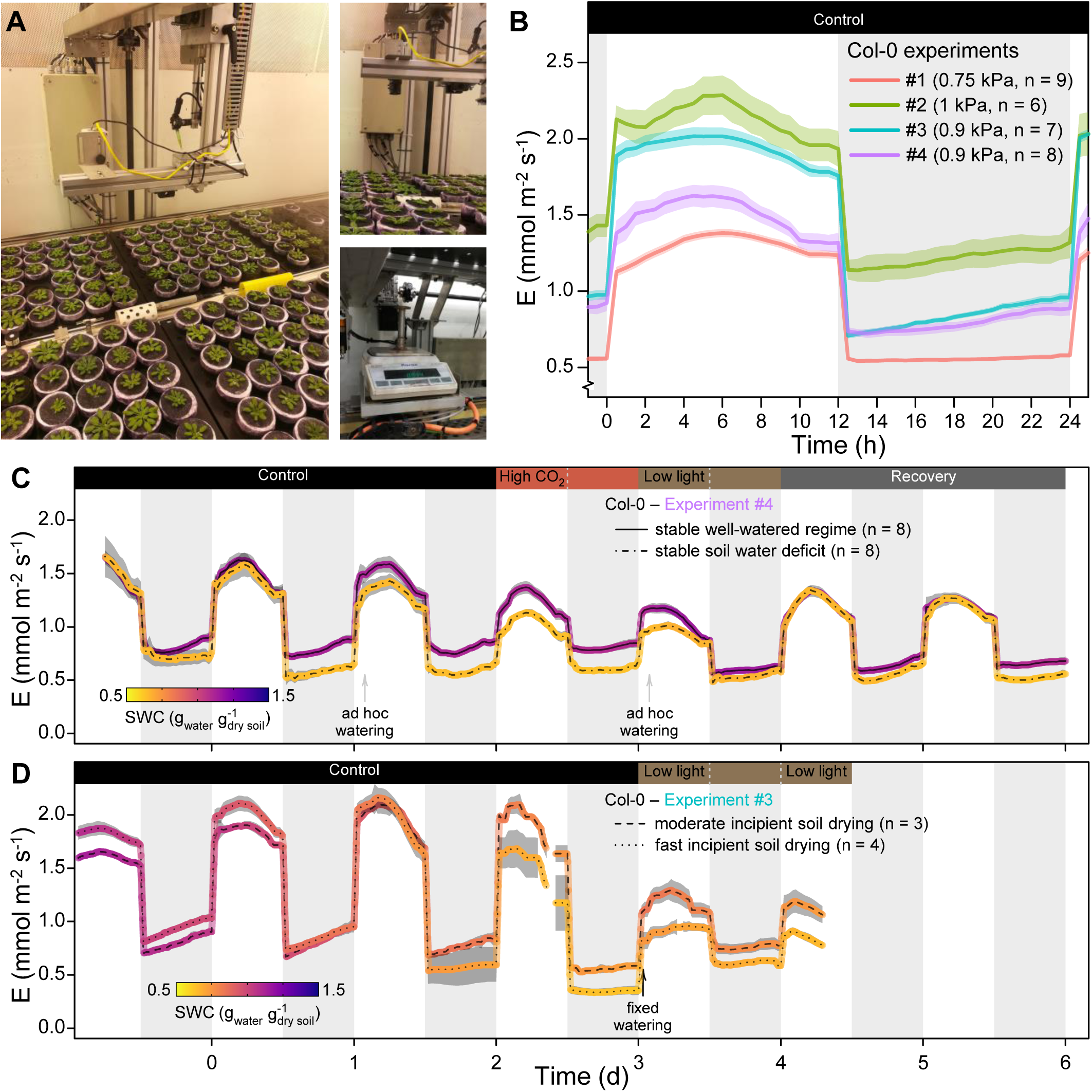
Using the PhenoLeaks phenotyping pipeline to analyse the diel transpiration in the wild type Col-0. **(A)** Overview of the Phenopsis chamber showing the automaton and the plants with soil covered with cellophane to prevent evaporation as well as a black grid to facilitate plant detection through image analysis (left), close-up above the plant surface featuring the zenithal camera for leaf area determination (top right), and close- up below the pots on the underlying mobile scale (bottom right) that periodically collected the weight data used to feed the PhenoLeaks pipeline for transpiration analysis. **(B)** Diel dynamics of transpiration rate (denoted E) of the wild type Col-0 in different experiments. Expanded view of the first complete diel cycle in control conditions across four independent experiments with slightly different settings (Supplemental Table S1). The vapour pressure deficit (VPD, in kPa) and the sample size (n) of each experiment is given in parentheses. **(C)** and **(D)** Transpiration dynamics of Col-0 throughout consecutive diel cycles in varying environmental conditions, for experiment #4 with two different stable watering regimes **(C)** and for experiment #3 where the target irrigation could not be adequately maintained **(D)**. Transpiration is colour-coded for the average soil water content (SWC) of each condition. In experiment #4, there were 2.75 d of control conditions (400 ppm CO_2_, 180 µmol m^-2^ s^-1^ irradiance), 1 d of elevated CO_2_ (800 ppm in the daytime), 1 d of low irradiance (50 µmol m^-2^ s^-1^ in the daytime), and 2 d of recovery in control conditions. In experiment #3, there were 3.9 d of control conditions and 1.5 d of low irradiance (same values as for experiment #4). The white and grey background depict the day (12 h) and night (12 h) periods, respectively. The shaded areas around the mean lines represent the means ± SE.

We also tested the robustness of the transpiration dynamics during several consecutive diel cycles, as well as their responses to changes in the environment. In experiment #4, we monitored the plants across about three diel cycles in control, growth conditions (180 µmol m^-2^ s^-1^ irradiance, 400 ppm CO_2_, 0.9 kPa VPD), before subjecting them to one day of elevated CO_2_ (800 ppm in the daytime), followed by one day of low irradiance (50 µmol m^-2^ s^-1^), plus two diel cycles of recovery (Figure 2C). The whole kinetics displayed a decreasing trend of transpiration rate as the experiment progressed, consistent with a gradual waterproofing of the Arabidopsis rosette as stomata acquire their function and cuticle get less permeable with leaf development (Pantin et al., 2013c; Kane et al., 2020; Hõrak et al., 2021). Both elevated CO_2_ and low irradiance decreased the amplitude of diel transpiration, consistent with their inhibiting effects on stomatal opening (Shimazaki et al., 2007; Zhang et al., 2018). Nevertheless, only low light noticeably affected the diel dynamics, showing an early afternoon preclosure (Figure 2C). Finally, two days of recovery in control environment showed that diel transpiration retrieved its original dynamics, with a transient and slight enhancement of the endogenous movements on the first day. These results suggest that the endogenous dynamics are robust and may respond to environmental changes with a transient acclimation period.

This interpretation was also supported by experiments where soil water content varied. In experiment #4, *ad hoc* watering was performed throughout the assay to adjust soil water content to a target value in all pots individually. When plants were subjected to a constant soil water deficit established one week before the transpiration assay, rosette growth was reduced but there was no major effect on the transpiration dynamics beyond the slight reduction in the baseline (Figure 2C). Thus, in these conditions of stable, moderate water stress, the acclimated plants globally maintained transpiration per unit leaf area, and conserved their native endogenous dynamics. By contrast, in experiment #3, plants were given a fixed watering amount after several days of transpiration monitoring. The initial water reserve plus the re-watering were not sufficient to maintain the soil water content (SWC) at a well-watered level, and the soil dried as transpiration proceeded (Figure 2D). We classified *a posteriori* the severity of the incipient soil drying into slow, moderate or fast based on the minimal soil water content that was reached. Expectedly, this was essentially driven by initial differences in rosette area (Supplemental Figure S2A), with larger plants showing faster soil water depletion than smaller plants. The speed of incipient soil drying appeared to enhance the decreasing trend of transpiration with ageing, accelerate the afternoon preclosure and reduce nighttime preopening (Figure 2D). Partial rewatering then lifted the transpiration baseline, even though low irradiance was simultaneously applied. Thus, a nascent soil water drying had more marked effects on the transpiration dynamics than an established water stress, even when the latter was slightly more acute (Figure 2C-D).

Overall, these diel dynamics gained from PhenoLeaks were consistent with previous data obtained on Arabidopsis using low-throughput, gas exchange analysis (Lascève et al., 1997; Sun et al., 2019) and stomatal aperture measurements (Li et al., 2020). Thus, our phenotyping pipeline reliably captured the broad patterns of transpiration throughout the diel cycle. Endogenous stomatal preclosure during the afternoon and preopening at night emphasized a diel coordination between days and nights, which is known to be under circadian control but may also be modulated by other rhythmic internal cues such as starch metabolism.

### Profiling the diel transpiration dynamics in two extreme starch mutants across environmental conditions highlights an endogenous stomatal rhythm with flexible coordination between day and night

We then examined two extreme mutants that were blocked in either starch synthesis for the starch-free *pgm* or starch breakdown for the starch-excess *sex1*. On the one hand, *pgm* cannot synthesise starch in any cell-type due to ineffective chloroplastic phosphoglucomutase (Caspar et al., 1985; Streb and Zeeman, 2012; Figure 1). On the other hand, *sex1* is defective in the plastidial α-glucan water, dikinase (GWD1) that is responsible for the first step of glucan phosphorylation necessary for starch breakdown (Caspar et al., 1991; Yu et al., 2001; Ritte et al., 2002). In *sex1*, daytime starch synthesis largely exceeds the residual nighttime degradation, resulting in a severe starch-excess phenotype (Streb and Zeeman, 2012; Figure 1A-B), which can be observed in mesophyll cells and also in the guard cells (Azoulay-Shemer et al., 2018; Figure 1C-D). Both *pgm* and *sex1* mutations result in very low starch turnover and generate similar diel variations in the carbon status: during the day, photosynthesis-derived sugars accumulate massively compared to the wild type, but then at night the absence or unusability of starch triggers a rapid depletion of available sugars, resulting in carbon starvation until dawn (Streb and Zeeman, 2012).

Here, the transpiration patterns of the starch mutants differed from that of the wild type, with *pgm* being more affected than *sex1*, as could be observed in experiment #4 that was then our reference experiment (Figure 3). For convenience, the first complete diel cycle is expanded in Figure 3A. To facilitate comparisons of the transpiration patterns, we extracted several parameters, where ‘E’ denotes a transpiration baseline while ‘A’ refers to an amplitude of variation in transpiration. To account for the global decrease of transpiration with ageing, the data were detrended before analysis (Methods). We quantified the mean transpiration over 24-h (E_diel_) and over day and night periods (E_day_, E_night_), as well as a set of descriptive parameters of the dynamics (Figure 4; Methods). While the mean transpiration rate over each period and the maximal diel amplitude (A_diel_) were similar in *pgm* as in Col-0 (Supplemental Figure S3A-D), their transpiration patterns were contrasted. At dawn, the rapid increase in transpiration due to stomatal opening observed at 30 min after the night-to-day transition (A_rapid op_) was reduced in *pgm* (Figure 4B), confirming previous findings (Lascève et al., 1997). Stomatal opening lasted longer in the mutant, which needed about 1 h more than Col-0 to reach the maximal transpiration (t_day max_, Figure 4D). This delay was associated with a different endogenous rhythm that appeared to propagate over the rest of the period. This was reflected by a highly significant difference in the rate of endogenous change of transpiration across the day (σ_day_, Supplemental Figure S3E) and a reduced stomatal preclosure in the afternoon (lower Σ_preclo_, Figure 4F). Overall, compared to the wild type, *pgm* showed less water loss in the morning but more water loss in the afternoon, so that E_day_ was similar in both genotypes. Similar conclusions could be drawn in the nighttime, when the period of slow stomatal closure after the rapid transition was prolonged in *pgm*: the time when transpiration reached its minimum during the night (t_night min_) was delayed by about 4 h (Figure 4E) in *pgm* compared to Col-0 in spite of similar amplitude of rapid reduction in transpiration due to stomatal closure at 30 min after the day-to-night transition (A_rapid clo_, Figure 4C). The highly significant difference in the rate of endogenous change of transpiration across the night (σ_night_, Supplemental Figure S3F) mirrored that observed in the daytime, and virtually no endogenous preopening was observed in *pgm* (Σ_preop_, Figure 4G), consistent with previous results (Lascève et al., 1997).

**Figure 3.**
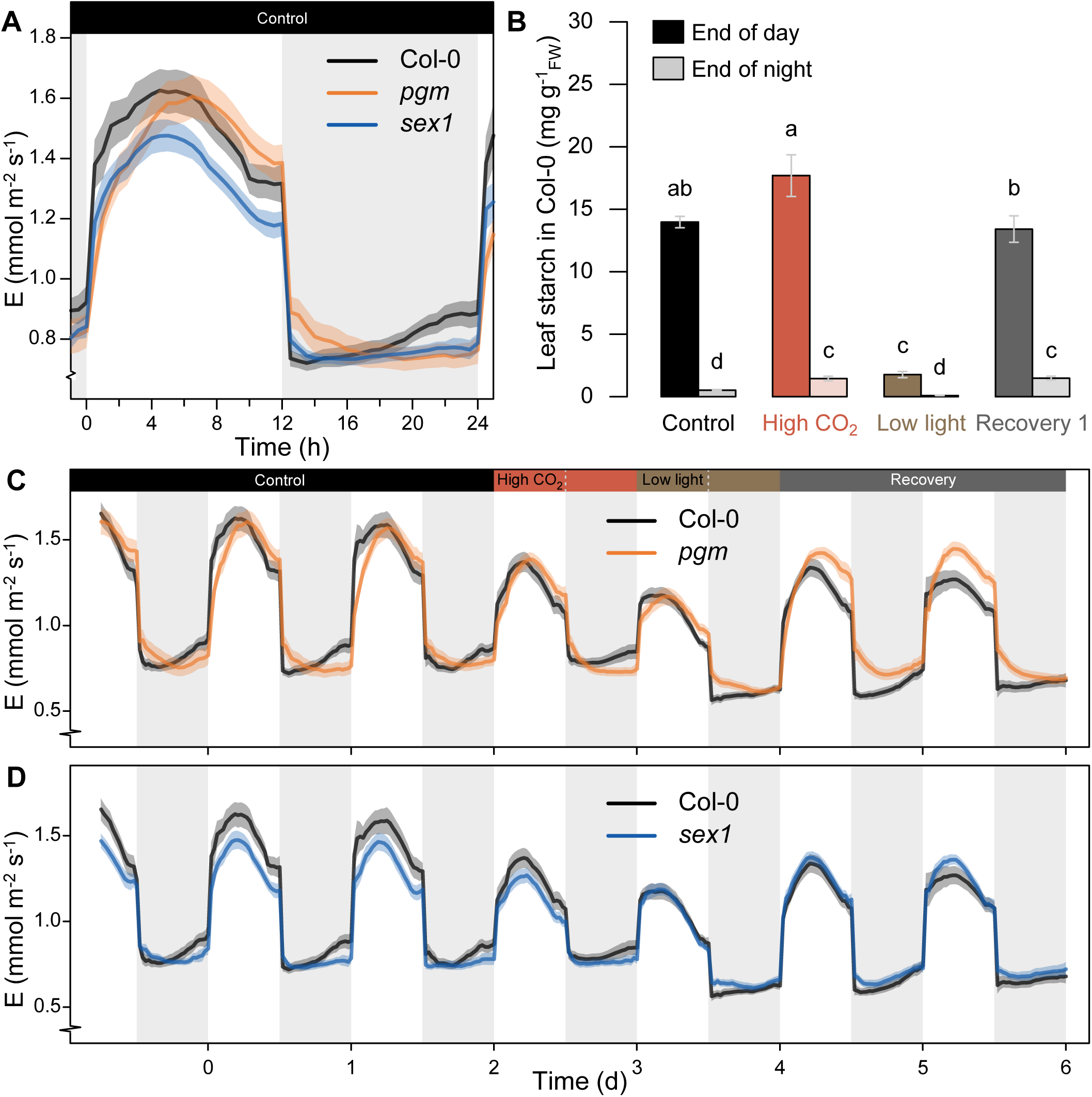
Analysis of diel transpiration of two extreme starch mutants, *pgm* and *sex1*. **(A)** Diel dynamics of transpiration rate of the wild type Col-0 compared to the starch-free *pgm* and the starch-excess *sex1*: expanded view of the first complete diel cycle under control conditions (experiment #4). **(B)** Leaf starch content determined at the end of the day and night periods in Col-0 for different environmental conditions during the kinetics hereinafter. Error bars are means ± SE. Letters denote significant differences after a Kruskal-Wallis test (α = 5%) followed by multiple comparisons of ranks. **(C)** and **(D)** Diel dynamics of transpiration rate of Col-0 compared to the starch-free *pgm* **(C)** and the starch-excess *sex1* **(D)** over consecutive diel cycles in the same conditions as in Figure 2C (experiment #4 – note that the data presented for the wild type are the same as in Figure 2C, stable well-watered regime). The colour-shaded areas around the mean lines represent the means ± SE.

**Figure 4.**
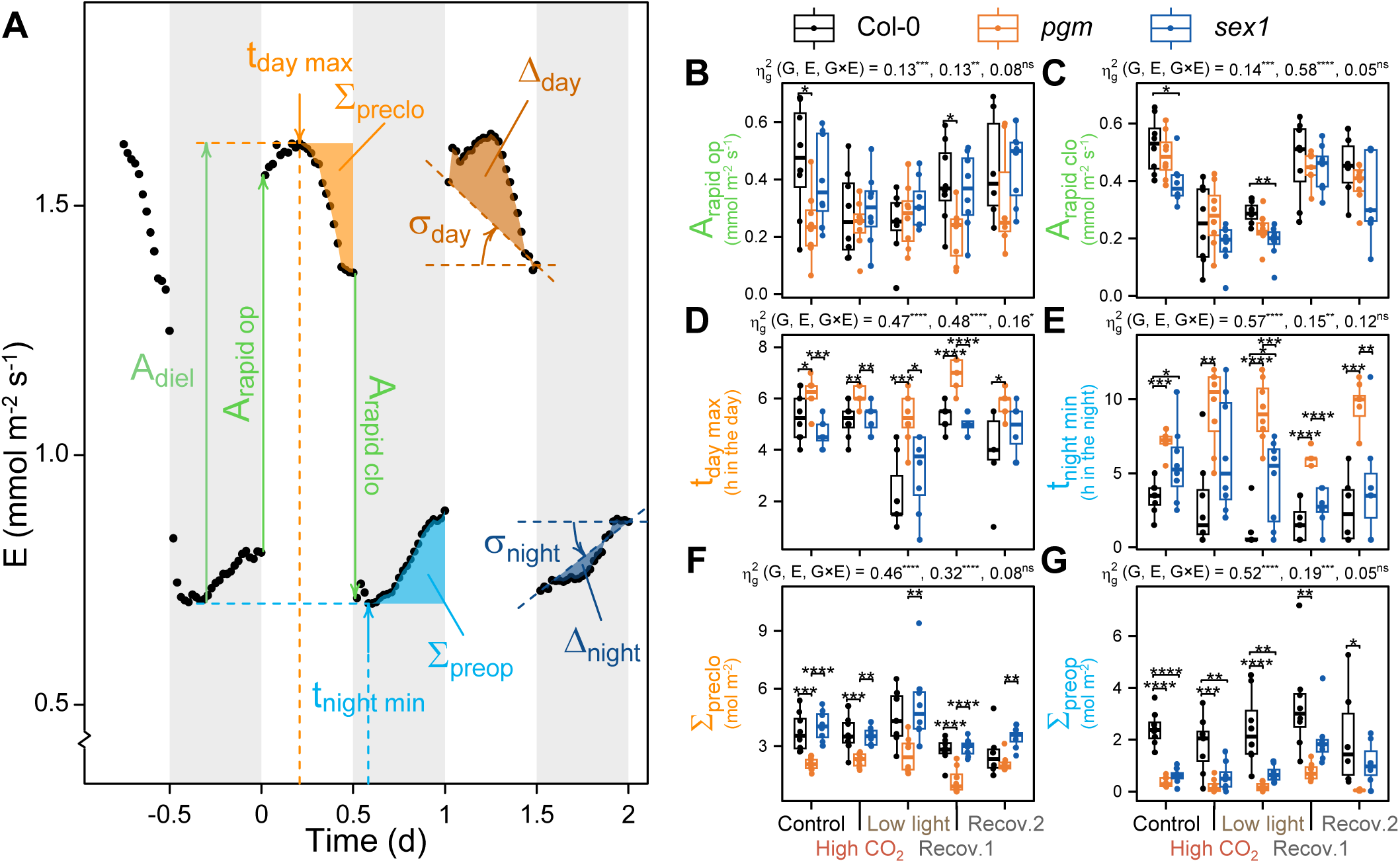
Quantitative analysis of the transpiration dynamics in Col-0, *pgm* and *sex1*. **(A)** Graphical example of the extraction of parameters on measured data, based on an individual plant of Col-0 in control conditions. For clarity, different parameters have been depicted on different days, but the initial 2.5 d were first averaged into a single 24-h kinetics, from which the parameters were then extracted. The data were linearly detrended before quantitative analysis. The parameters are detailed in the Methods section. **(B)** to **(G)** Boxplots of selected parameters extracted from the measured data for the different genotypes and environments: **(B)** rapid stomatal opening at the day-to- night transition (A_rapid op_), **(C)** rapid stomatal closure at the night-to-day transition (A_rapid clo_), **(D)** time of maximal daytime transpiration (t_day max_), **(E)** time of minimal nighttime transpiration (t_night min_), **(F)** cumulative afternoon preclosure after t_day max_ (Σ_preclo_) and **(G)** cumulative nighttime preopening after t_night min_ (Σ_preop_). For each parameter, a two-way ANOVA was done and the effect sizes (η^2^_g_) of the genotype (G), of the environment (E) and of their interaction (G×E), as well as their respective significance level, are reported above each corresponding plot. Pairwise t-tests were systematically done to assess the differences between genotypes in a given environment, and the differences are reported on the plot when significant. Significance codes are as follows: ^ns^, not significant; *, 0.05 ≤ p_val_ < 10^-2^; **, 10^-2^ ≤ p_val_ < 10^-3^; ***, 10^-3^ ≤ p_val_ < 10^-4^; ****, 10-4 ≤ pval.

These data straightaway discard a univocal, instantaneous role of soluble sugars in driving stomatal movements at any time of the diel cycle. Let us assume that sugars promote stomatal opening, as demonstrated with the double mutant *stp1 stp4* that lacks key glucose importers in the guard cells and barely opens stomata after 2-h illumination (Flütsch et al., 2020a). A lack of starch-derived sugars in *pgm* during the night or at the night-to-day transition may result in slower stomatal (pre)opening (Figure 4B,G). However, shortly after the onset of illumination, massive amounts of soluble sugars are found in the leaves of *pgm* (Caspar et al., 1985). For instance, 2 h after dawn, the content in sucrose and reducing sugars is already more than two times higher in *pgm* than in wild-type leaves (Gibon et al., 2004b; Pal et al., 2013), while transpiration of the mutant is still lagging behind (Figure 3A). It is therefore unlikely that the lack of sugars impedes stomatal opening during several hours in *pgm*. Alternatively, the idea that sugar excess in *pgm* would hamper stomatal opening in the morning is not either consistent with the observation that in the afternoon, sugar content remains higher in the mutant than in Col-0, while its stomata start closing later.

Analysing *sex1* further confirmed the absence of univocal relationship between sugar content and diel stomatal movements. In this mutant, the mean transpiration rates, the diel amplitude, the rapid changes at the transitions, and the daytime pattern were essentially similar as in Col-0 (Figure 4B-D,F, Supplemental Figure S3A-E,G), by contrast with the nighttime endogenous rhythm. Nighttime transpiration after the light-to-dark transition was almost constant in *sex1* (σ_night_ close to zero, Supplemental Figure S3F), making it difficult to estimate t_night min_ with good precision for this mutant (high variance of this parameter, Figure 4E). Nevertheless, nighttime preopening was clearly reduced in *sex1*, resembling *pgm* (Figure 4G). Thus, while *pgm* and *sex1* have low starch turnover and broadly similar sugar content throughout the diel cycle (Caspar et al., 1985, 1991), their transpiration phenotypes appeared to be shared only at night. This suggests that starch and sugar metabolisms do not directly drive the diel stomatal movements, but bring flexibility in the regulation of the endogenous rhythm that coordinates daytime and nighttime stomatal behaviour.

It was also clear from the environmental manipulations that variations in starch turnover are not associated with coherent changes in the diel dynamics of transpiration. The environmental scenario in experiment #4 was designed to induce variations in starch turnover, which was achieved in Col-0 (Figure 3B). Doubling the atmospheric CO_2_ (800 ppm) in the daytime enhanced end-of-day starch accumulation by 25%, while the remaining starch at the end of the night almost tripled, but this was not associated with remarkable changes in the endogenous stomatal rhythm (Figures 3 and 4, Supplemental Figure S3). A subsequent day at low irradiance (50 µmol m^-2^ s^-1^) dramatically reduced starch synthesis and turnover, but it did not phenocopy the effects of the *pgm* or *sex1* mutations, even triggering opposite effects with t_day max_ being advanced by almost 3 h (Figures 3 and 4, Supplemental Figure S3). Thus, manipulating the environment affected the diel transpiration pattern in Col-0, but not as would be expected from the changes in starch turnover. Most importantly, *pgm* and *sex1* responded to the environment essentially as the wild type (Figure 3C,D). This observation was supported by the ANOVA effect sizes (η^2^, Figure 4B-C, Supplemental Figure S3), which remained always lower for the interaction than for the main effect(s) of the genotype or the daytime environment. This result further supports the idea that changes in starch turnover are not required to support the changes in transpiration dynamics induced by the environments.

These conclusions were also verified in experiment #3 that comprised several starch mutants shared with experiment #4, including *pgm* and *sex1*. The transpiration kinetics (Supplemental Figure S4A) resembled that obtained in experiment #4 (Figure 3C-D). In control conditions, we confirmed the delayed opening and preclosure of *pgm* in the daytime (Supplemental Figure S4C-D), and the reduced nighttime preopening of *pgm* and *sex1* (Supplemental Figure S4E-F). We also confirmed that low light changes the daytime transpiration pattern in a different way (Supplemental Figure S4A,C-D), especially in the mutants that were barely affected by the variations in soil water content because they were smaller than Col-0 (Supplemental Figure S2A,D).

To summarize, starch turnover *per se* or instantaneous sugar content are likely not instrumental in the endogenous rhythm of transpiration. Nonetheless, severe mutations in starch metabolism or low irradiance interfere with the endogenous stomatal rhythm in various ways at different times of the diel cycle, revealing a flexible coordination between daytime and nighttime dynamics.

### Harmonic modelling of the endogenous transpiration rhythm captures the flexible coordination between daytime and nighttime stomatal movements

The flexible coordination between day and night patterns of transpiration, and the circadian nature of the endogenous stomatal rhythm (Hubbard and Webb, 2015), prompted us to build an overarching framework that could capture daytime and nighttime traits through a common set of parameters. This was achieved by designing a sinusoidal model (Figure 5; Methods), where diel transpiration shows step changes at the day/night transitions, and oscillates throughout the whole day and night cycle according to a periodic, sinusoidal signal. The temporal response of stomata to a rapid change in light intensity, which predominates at the day/night transitions, follows an exponential kinetics with a time constant (*i*.*e*. the duration for reaching 63% of the full response) typically of about 20 min for opening and 5 min for closure in Arabidopsis (Matthews et al., 2018). Such temporal variations were too rapid to be captured with good precision in our experiments due to the 30-min resolution, so we used a square wave at the day/night transitions (equivalent to a time constant of 0 min) in the model. Hence, stomatal opening and closure at the day/night transitions were considered as broadly symmetrical, which was a reasonable assumption given that similar values were obtained for A_rapid op_ and A_rapid clo_ in steady conditions (Figure 4B-C). For the rest of the diel cycle, we implemented a periodic signal composed of a fundamental sine wave (period fixed to 24 h) and its second harmonic (period fixed to 12 h), which allowed for asymmetry between daytime and nighttime patterns. Note that such sinusoidal signal may convey different behaviours of the circadian oscillator in the light and dark, but may also account for endogenous diel rhythms that are partly independent from the clock, like starch metabolism or leaf water status, introducing additional controls at different times of the diel cycle.

**Figure 5.**
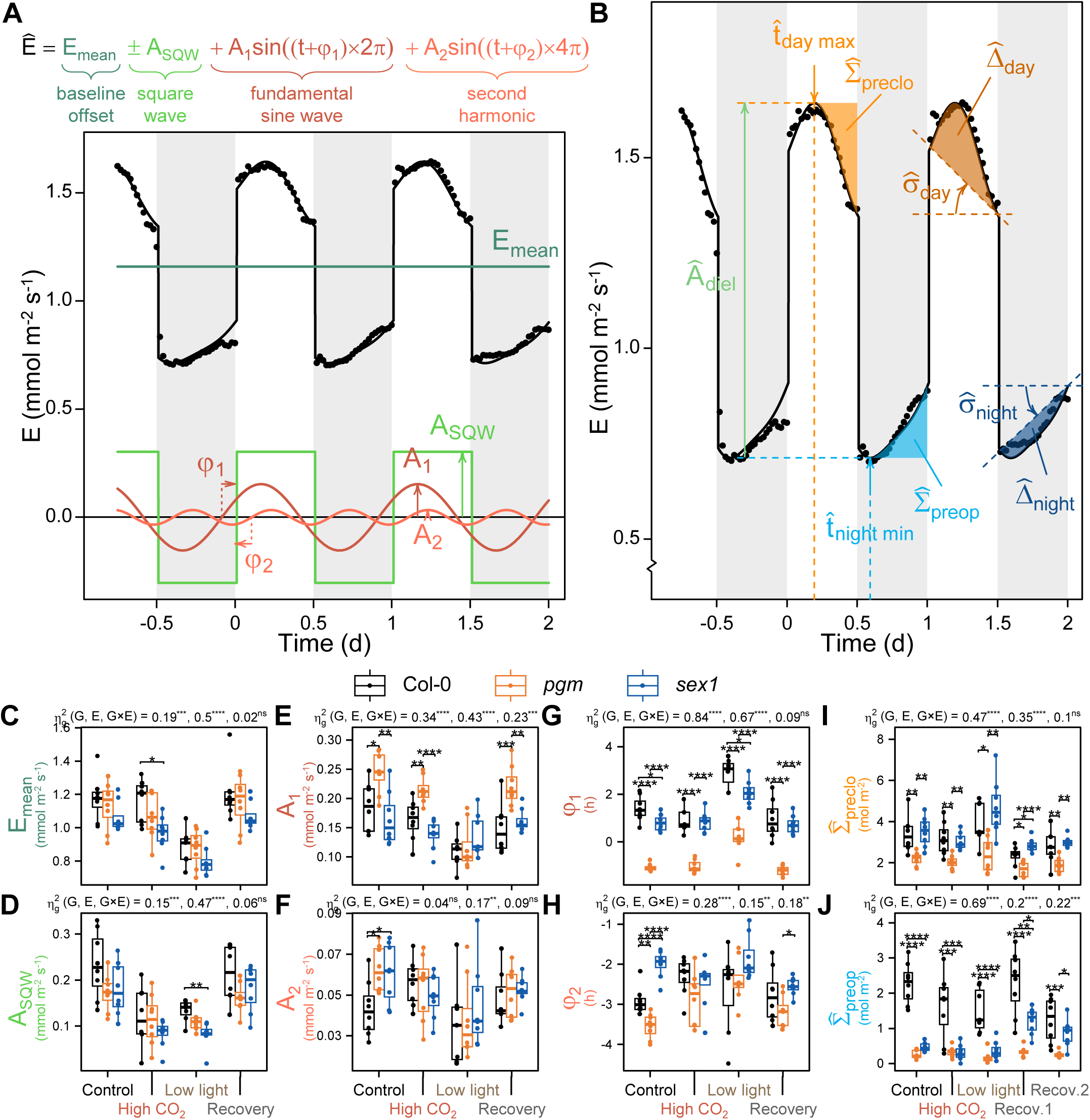
Modelling diel transpiration dynamics and obtaining associated parameters in Col-0, *pgm and sex1*. **(A)** Structure of the sinusoidal model, graphical example of the resulting curve (E, black line) and illustration of each component (coloured lines) using the set of parameters fitted to the data (black points) of the same Col-0 individual plant as shown in Figure 4A. **(B)** Graphical example of the extraction of quantitative parameters on fitted data, similar as for Figure 4A, using the same individual plant. **(C)** to **(H)** Boxplots of fitted parameters for the different genotypes and environments: **(C)** Mean transpiration rate over a 24 h period (E_mean_), **(D)** semi-amplitude of the square wave (A_SQW_), **(E)** semi-amplitude of the fundamental sine wave (A_1_), **(F)** semi-amplitude of the second harmonic (A_2_), **(G)** phase of the fundamental sine wave (φ_1_) and **(H)** phase of the second harmonic (φ_2_). **(I)** and **(J)** Boxplots of selected parameters extracted from the fitted data for the different genotypes and environments: **(I)** cumulative afternoon preclosure (Σ_preclo_) and **(J)** cumulative nighttime preopening (Σ_preop_). Significance codes for ANOVA effect sizes (Σ^2^_g_) and pairwise t-tests are like those in Figure 4.

The core model had only six parameters that were shared between day and night and easily interpretable in analogy with sinusoidal signal analysis (Figure 5). The ‘baseline offset’ corresponded to the mean transpiration rate over a 24 h period (E_mean_, mmol m^-2^ s^-1^) and was therefore mathematically almost identical to E_diel_. The semi-amplitude of the square wave (A_SQW_, mmol m^-2^ s^-1^) accounted for stomatal opening and closure at the day/night transitions, representing rapid stomatal responses similar to A_rapid op_ and A_rapid clo_. Each harmonic was characterized by its the semi-amplitude (A_1_ for the fundamental wave and A_2_ for the second harmonic, mmol m^-2^ s^-1^), setting the magnitude of variation in the endogenous rhythm around the baseline, and by its phase (ϕ_1_ and ϕ_2_, h), setting the timing of the endogenous rhythm (advance or delay compared to the reference square wave). This model could simply be fitted using linear regression once appropriately transformed (Methods). Using experiment #4, we initially fitted one set of parameters for each plant and for each diel cycle separately, except the period in control conditions where the diel cycles were fitted jointly. An example for Col-0 in control conditions is presented in Figure 5A, showing excellent agreement between the crude model and the data. Similar quality was obtained for the other genotypes and environmental conditions, as illustrated by different examples in Supplemental Figures S5, S6 and S7. Nevertheless, some plants showed an enhanced decreasing trend upon switching to low light, which appeared to propagate over the whole diel cycle and was not properly captured by the model (*e*.*g*. S5A, S6B, S7B). We interpreted this trend as a transient acclimation to the new environmental conditions. This interpretation also agreed with the enhanced endogenous rhythm of the first recovery cycle compared to the second one, as well as the stronger response to the incipient soil drying compared to the stable water deficit (Figure 2C-D). We therefore augmented the model with an ‘acclimation pulse’, adding a linear effect of slope α (mmol m^-2^ s^-1^ d^-1^) for 24 h when plants are subjected to a change in the environment. This additional parameter improved the fit quality especially for the low light period, and allowed us to use one common set of parameters for both recovery cycles, with only the first one being affected by the acclimation pulse (Supplemental Figures S5, S6, S7). Across the whole experiment #4 (19 conditions × 4 environmental periods × 8 replicates), the RMSE averaged 0.031 mmol m^-2^ s^-1^. It was then straightforward to extract further parameters from the fitted curves following a homogeneous process as with the observed ones (Figure 5B).

### Blocking starch metabolism through severe mutations induces a phase shift in the endogenous stomatal rhythm that is not phenocopied upon low light in the wild type

Analysing the fitted parameters highlighted the architecture of the diel dynamics, and revealed a phase shift in the starch mutants (Figure 5C-H). In control conditions and in all genotypes, the magnitude of the rapid responses at the day/night transitions (A_SQW_) was similar as the magnitude of the endogenous rhythm (2×[A_1_+A_2_]), with the fundamental wave largely dominating over the second harmonic (A_1_ being three to four times higher than A_2_). Accordingly, the fundamental sine peaked around the middle of the daytime, about 1.4 h before midday for Col-0 in control conditions (ϕ_1_ = +1.41 ± 0.19 h, meaning that the peak is at 24 / 4 – 1.41 = 4.59 h after the light switches on). The second harmonic peaked around midday too, and thus also around midnight (for Col-0, ϕ_2_ = −2.91 ± 0.13 h, meaning that the peaks are at 12 / 4 + 2.91 = 5.91 h after the light switches on and off). Consequently, the variations of the second harmonic were relatively well synced with that of the fundamental sine during the day (thereby enhancing the endogenous transpiration rhythm), but had an opposite behaviour at night (thereby attenuating the endogenous rhythm). Manipulating the environment reduced the amplitudes, with A_1_ being more sensitive to low irradiance, while A_2_ remained mostly unaffected. The environment also affected the acclimation pulse, especially low light, and this response was shared by all genotypes (Supplemental Figure S8). By contrast, the genotypes had strong effects over the phases. We uncovered a marked phase shift on ϕ_1_ with more than 2 h delay in *pgm*, which was conserved across the environments (Figure 5G) and even under stable soil water deficit (Supplemental Figures S2C, S9). We observed a smaller phase delay of 45 min for ϕ_2_. In *sex1*, ϕ_1_ was slightly delayed by 30 min, while ϕ_2_ was 1 h ahead compared to Col-0. On the other hand, low irradiance advanced ϕ_1_ by almost 1.5 h in all genotypes. Thus, shifts in the phase of the fundamental sine resulted in a global advance (low light) or delay (*pgm* and to a lesser extent *sex1*) in the endogenous rhythm, while independent phase modulation of the second harmonics resulted in different effects on the afternoon preclosure (*pgm* only) and nighttime preopening (*pgm* and *sex1*), which were recapitulated by the model (compare Figures 3E,G and 4I-J).

These conclusions were corroborated by experiment #3. There were differences between both experiments in the range of the fitted parameters (Supplemental Figure S10), reflecting differences in the experimental data (Figure 3C-D, Supplemental Figures S4). However, in control conditions, the genotypic differences were widely conserved, including the striking phase shift in *pgm*, and to a lesser extent *sex1* (Supplemental Figure S10D). Under low light, results of the wild type were flawed by its lower soil water content (Figure 2D). Nonetheless, in *pgm* and *sex1* plants that were smaller and thus not affected by soil drying (Supplemental Figure S2A), low light triggered an advance in ϕ_1_, confirming the results of Experiment #4 – even though low light in the latter was preceded by one day at high CO_2_. Overall, the major effects of the starch mutants were conserved across experiments #3 and #4.

### Comparing mutants of starch metabolism pinpoints a key role for maltose export in controlling the endogenous rhythm of transpiration

To get deeper insight into the connection between diel transpiration dynamics and starch metabolism, we then scrutinized the behaviour of the individual mutants, starting by the biosynthesis pathway downstream of PGM. We first examined two mutants with mild reduction in starch synthesis, which both exhibited differences in transpiration dynamics compared to the wild type, yet not on the same parameters (Figure 6, Supplemental Figure S11). On the one hand, *isa1* is impaired in the debranching enzyme isoamylase 1, which contributes to the structure of starch; it accumulates phytoglycogen rather than only starch and shows premature exhaustion of its reserves during the night (Delatte et al., 2005; Wattebled et al., 2005; Figure 1). Here, *isa1* had a transpiration phenotype resembling that of *pgm*, with disturbed rates of change of the endogenous rhythm (σ_day_ and σ_night_, Supplemental Figure S11G,K) and reduced nighttime preopening (Σ_preop_, Figure 6J) that coincided with a ϕ_1_ delay of 1.5 h (Figure 6G). On the other hand, *ss4* is affected in starch synthase 4, which is required for the initiation of starch granules; it has reduced starch synthesis and, contrary to *isa1*, incomplete nighttime breakdown (Roldán et al., 2007; Crumpton-Taylor et al., 2013; Lu et al., 2018; Figure 1). In our assay, the endogenous transpiration pattern of *ss4* showed reduced A_diel_ (Supplemental Figure S11B) but unaltered phases. These results confirmed that disturbing starch metabolism may trigger different changes in the diel dynamics of transpiration, but that starch turnover does not simply drive the endogenous stomatal rhythm.

**Figure 6.**
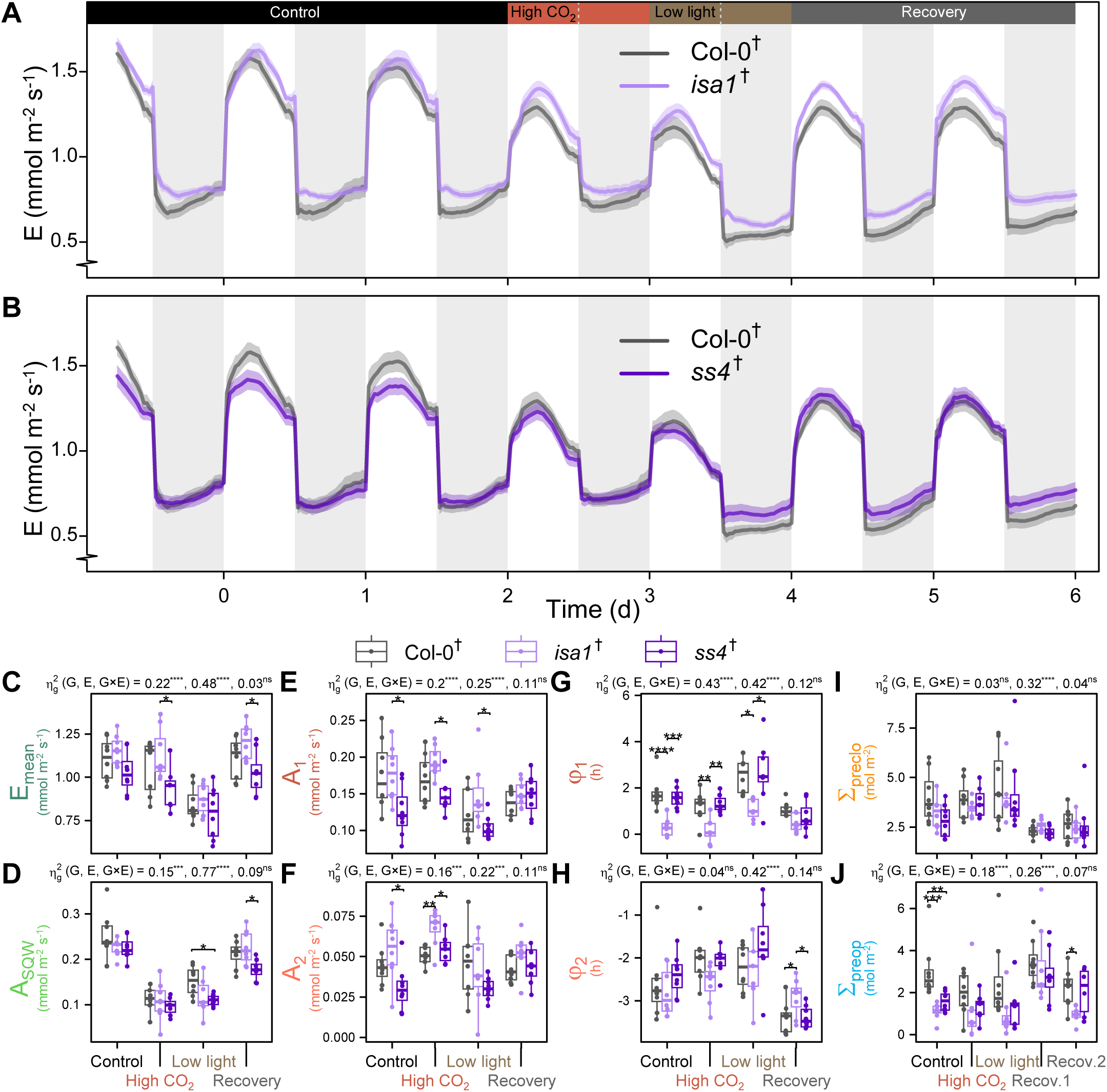
Analysis of diel transpiration of *isa1* and *ss4*, two mutants of the starch biosynthesis pathway. **(A)** and **(B)** Diel dynamics of transpiration rate of the wild type Col-0 compared to *isa1* **(A)** and *ss4* **(B)** in the same conditions as in Figure 3. The colour-shaded areas around the mean lines represent the means ± SE. **(C)** to **(H)** Boxplots of fitted parameters for the different genotypes and environments: **(C)** E_mean_, **(D)** A_SQW_, **(E)** A_1_, **(F)** A_2_, **(G)** ϕ_1_ and **(H)** ϕ_2_. **(I)** and **(J)** Boxplots of selected parameters extracted from the measured data for the different genotypes and environments: **(I)** Σ_preclo_ and **(J)** Σ_preop_. Abbreviations for parameters and significance codes for ANOVA effect sizes (η^2^_g_) and pairwise t-tests are like those in Figures 4 and 5. The dagger in Col-0^†^, *ss4*^†^ and *isa1*^†^ indicates that these lines harbour a luciferase reporter that was not used in this study (Methods).

We next examined the influence of starch degradation downstream of SEX1 and uncovered a key role for chloroplastic maltose transport in the diel transpiration dynamics (Figure 7, Supplemental Figure S12). Maltose is the main product of nighttime starch breakdown in the mesophyll chloroplasts; it is generated by β-amylases and then exported to the cytosol by the MALTOSE EXCESS 1 (MEX1) chloroplastic transporter (Niittylä et al., 2004; Weise et al., 2004, 2005; Figure 1A). The *mex1* mutant over-accumulates maltose in the chloroplast and is impaired in starch breakdown (Niittylä et al., 2004; Lu et al., 2006). The function of MEX1 in starch metabolism of the guard cells is not elucidated, yet. In our basic observations, the guard cells of *mex1* appeared essentially devoid of starch at the end of the night and day, contrary to the rest of the leaf that had starch in excess (Figure 1B-D). This spatial pattern was unique across all the mutants we monitored. More thorough observations by Flütsch (2020) have shown that *mex1* guard cells do not over-accumulate starch at the end of the night, but do not degrade starch within 1 h of light, suggesting MEX1 is not the sole exporter of starch degradation products in the guard cells. In control conditions, the endogenous transpiration dynamics of *mex1* showed disorders like in *pgm* and *isa1*, with altered σ_day_ and σ_night_ (Supplemental Figure S12G,K) and reduced nighttime preopening (Figure 7K) that were subtended by a ϕ_1_ delay of 2.1 h. Moreover, *mex1* was exquisitely sensitive to changing environmental conditions, especially low irradiance. The baseline transpiration E_mean_ was much more affected by elevated CO_2_ and low irradiance in *mex1* than in the wild type, yielding highly significant differences between both genotypes in these non-control conditions (Figure 7D). Both E_day_ and E_night_ were affected (Supplemental Figure S12E,I). These aberrant stomatal responses might arise from a loss of chloroplast integrity in *mex1*, which develops chlorotic leaves with chloroplasts showing autophagy-like degradation (Stettler et al., 2009). For instance, chloroplast degradation could disrupt the production of regulatory signals from the mesophyll (Lawson and Matthews, 2020), or prevent turgor generation within the guard cells (Azoulay-Shemer et al., 2015). However, disturbed environmental conditions were unlikely to produce irreversible chloroplastic damages: upon return to the original growth conditions, the diel dynamics remarkably recovered its original endogenous rhythm and the same baseline as the wild type (Figure 7, Supplemental Figure S12). Across our experiment, no mutant other than *mex1* showed such marked sensitivity to environmental conditions, pinpointing the importance of the transporter therein.

**Figure 7.**
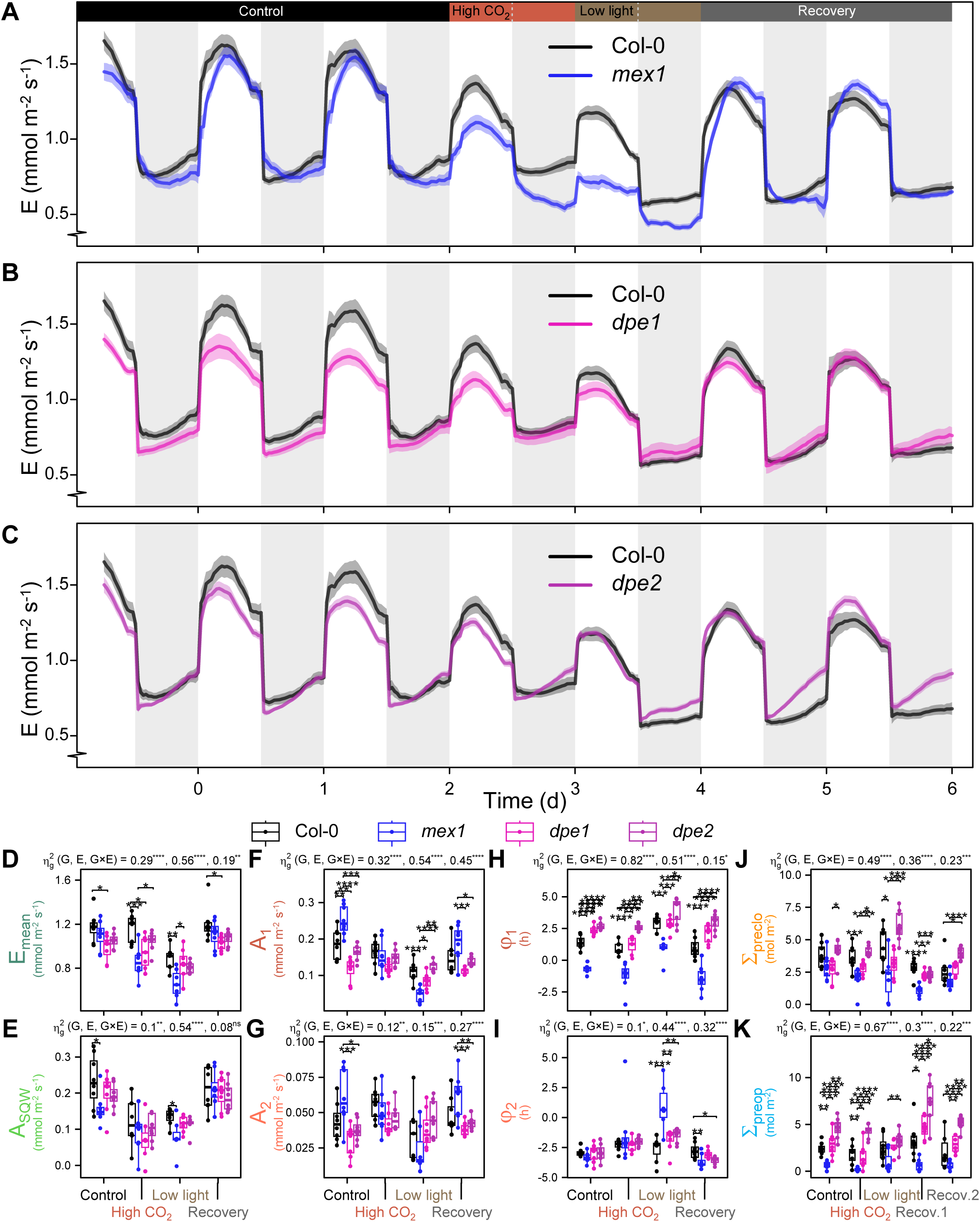
Analysis of diel transpiration of *mex1, dpe1 and dpe2*, three mutants of the starch breakdown pathway. **(A)** to **(C)** Diel dynamics of transpiration rate of the wild type Col-0 compared to *mex1* **(A)**, *dpe1* **(B)** and *dpe2* **(C)** in the same conditions as in Figure 3. The colour-shaded areas around the mean lines represent the means ± SE. **(D)** to **(I)** Boxplots of fitted parameters: **(D)** E_mean_, **(E)** A_SQW_, **(F)** A_1_, **(G)** A_2_, **(H)** φ_1_ and **(I)** φ_2_. **(J)** and **(K)** Boxplots of selected parameters extracted from the measured data: **(J)** Σ_preclo_ and **(K)** Σ_preop_. Abbreviations for parameters and significance codes for ANOVA effect sizes (Σ^2^_g_) and pairwise t-tests are like those in Figures 4 and 5.

Analysis of two other mutants in this pathway further supported the view that maltose export influences the diel dynamics of transpiration (Figure 7, Supplemental Figure S12). Next to maltose, glucose is produced by the chloroplastic disproportionating enzyme 1 (DPE1) during starch breakdown (Critchley et al., 2001), and is further exported to the cytosol by the glucose transporter pGlcT (Cho et al., 2011; Figure 1A). The glucose route is parallel to the maltose route and is thought to be minor (Streb and Zeeman, 2012). Here, blocking the glucose pathway in *dpe1* affected starch turnover similarly as in *mex1* (Figure 1B), but resulted in contrasting transpiration dynamics: the diel pattern of *dpe1* (Figure 7C) resembled more that of the wild type, yet ϕ_1_ was advanced by 1 h and t_night min_ by 2.4 h, while A_1_ was reduced (Figure 7F,H, Supplemental Figure S12J). Contrary to *mex1*, no major difference was observed upon changing the environmental conditions. We also looked at *dpe2*, a mutant affected in the cytosolic disproportionating enzyme 2 that uses the maltose exported by MEX1 to release glucose (Streb and Zeeman, 2012; Figure 1A). The *dpe2* mutant has a starch-excess phenotype (Figure 1B), and accumulates extremely high levels of maltose in the cytosol throughout the diel cycle (Lu et al., 2006; Chia et al., 2004; Lu and Sharkey, 2004; Weise et al., 2005). We observed transpiration phenotypes in *dpe2* (Figure 7C) that were opposite to that in *mex1*, with enhanced nighttime preopening as well as lower t_night min_ and higher σ_night_, which were concomitant with a ϕ_1_ advance of 1.3 h compared to Col-0 in control conditions (Figure 7H,K, Supplemental Figure S12J,K). Switching to elevated CO_2_ or low irradiance triggered similar changes in *dpe2* as in the wild type, but return to the original conditions induced an upsurge in stomatal preopening (Figure 7C,K, Supplemental Figure S12K). These results supported a role for maltose export in promoting nighttime stomatal preopening by advancing the phase of the endogenous rhythm or gating its amplitude at night. However, the lack of knowledge about the involvement of maltose and MEX1 during starch breakdown in the guard cells (Santelia and Lunn, 2017) made it difficult to understand whether the observed phenotypes were due to guard-cell-autonomous metabolic dysfunctions, or remote effects from the mesophyll metabolism.

### Neither guard-cell-autonomous nor bulk-leaf starch turnover may directly sustain diel stomatal dynamics

We next analysed genotypes that are differentially affected in guard-cell *vs*. mesophyll-cell starch metabolism (Figure 8, Supplemental Figures S13, S14, S15). We focused on the *pgi* mutant, which is defective in the plastidial phosphoglucose isomerase that produces Glc6P from Fru6P upstream of PGM (Yu et al., 2000; Figure 1A). While starch synthesis is largely impaired in *pgi*, some cell types can bypass the PGI step by importing Glc6P directly from the cytosol (Overlach et al., 1993; Niewiadomski et al., 2005; Flütsch et al., 2022; Figure 1A). Accordingly, *pgi* is essentially starch-deficient in the mesophyll cells, but retains starch synthesis in several cell types including the guard cells (Tsai et al., 2009; Kunz et al., 2010; Azoulay-Shemer et al., 2016; Figure 1D), resulting in a reduced daytime starch accumulation when measured at the bulk-leaf level (Figure 1B-C). Here, no difference in the transpiration dynamics was found between *pgi* and the wild-type, except a slight shift in the baseline (Figure 8A,D-K, Supplemental Figure S13). Does this result mean that guard cells hold the pool of starch that autonomously supports stomatal dynamics?

**Figure 8.**
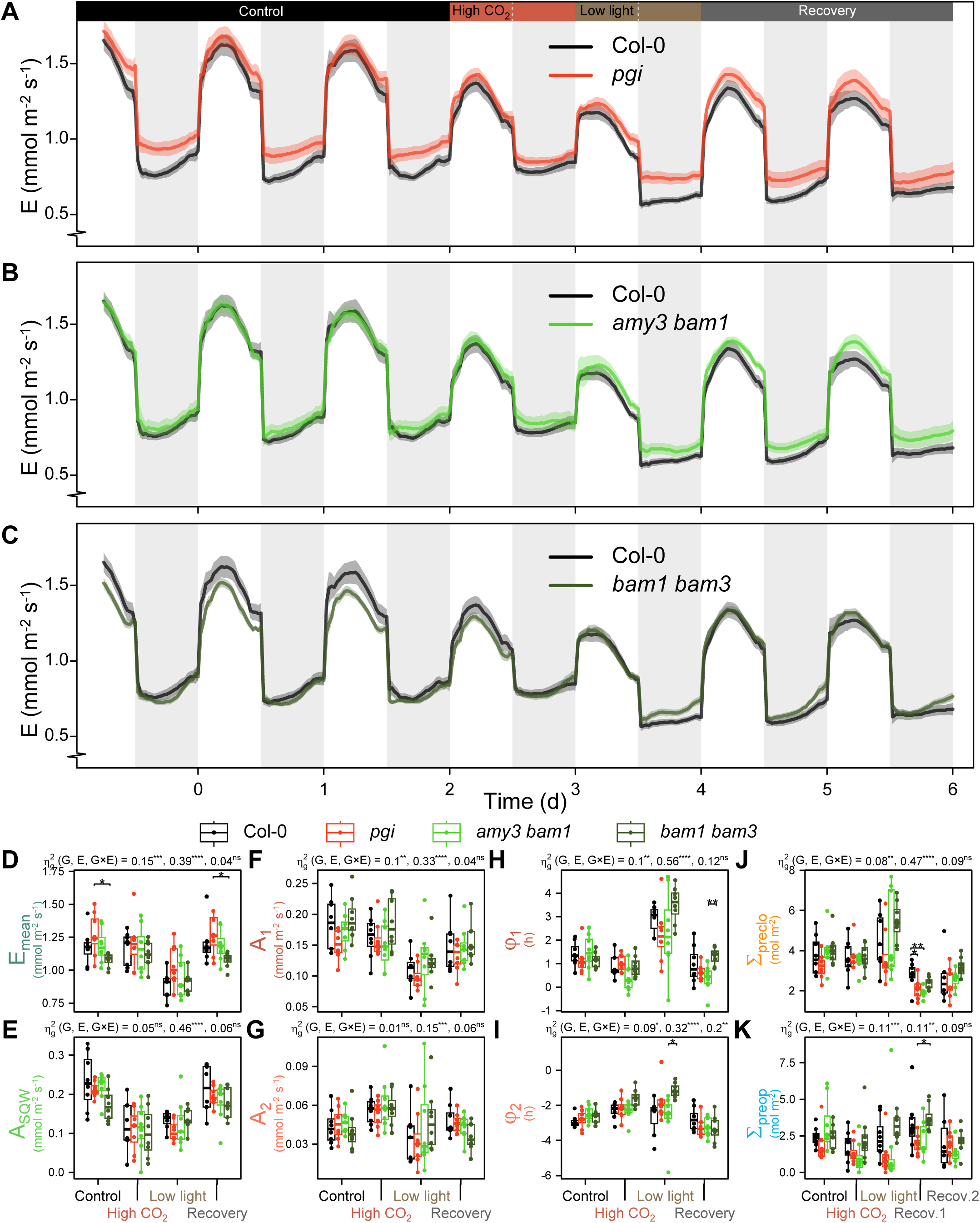
Analysis of diel transpiration of *pgi, amy3 bam1 and bam1 bam3*, three mutants with contrasting starch metabolism in different tissues. **(A)** to **(C)** Diel dynamics of transpiration rate of the wild type Col-0 compared to *pgi* **(A)**, *amy3 bam1* **(B)** and *bam1 bam3* **(C)** in the same conditions as in Figure 3. The colour-shaded areas around the mean lines represent the means ± SE. **(D)** to **(I)** Boxplots of fitted parameters: **(D)** E_mean_, **(E)** A_SQW_, **(F)** A_1_, **(G)** A_2_, **(H)** φ_1_ and **(I)** φ_2_. **(J)** and **(K)** Boxplots of selected parameters extracted from the measured data: **(J)** Σ_preclo_ and **(K)** Σ_preop_. Abbreviations for parameters and significance codes for ANOVA effect sizes (Σ^2^_g_) and pairwise t-tests are like those in Figures 4 and 5.

We tested this cell-autonomous hypothesis, but could not find evidence to support it. Starch degradation in the guard cells is achieved through the guard-cell specific and synergistic action of the isoforms α-amylase AMY3 and β-amylase BAM1 (Horrer et al., 2016; Flütsch et al., 2020b; Figure 1A). The double mutant *amy3 bam1* is severely impaired in starch degradation in the guard cells but has intact starch turnover in the bulk leaf (Figure 1B-D). Here, the diel dynamics of transpiration of *amy3 bam1* were virtually indistinguishable from that of Col-0 (Figure 8B), just like the single mutants (Supplemental Figure S14A-B). No significant difference could be detected in the measured or fitted parameters between Col-0 and *amy3 bam1* across all environments (Figure 8D-K, Supplemental Figure S13). This result demonstrated that guard cell-autonomous starch breakdown is not required to achieve proper diel stomatal movements.

Alternatively, it might be plausible that the pool of starch from the mesophyll can substitute for that from the guard cells in generating a control on stomata, and vice versa. We therefore looked at two other mutants affected in the family of β-amylases. BAM3 is normally the main isoform involved in mesophyll starch degradation but in its absence, BAM1, also active in guard cells, partly compensates its function (Fulton et al., 2008). Accordingly, the *bam3* single mutant has an intermediate starch-excess phenotype in the mesophyll only, while the *bam1 bam3* double mutant has a severe starch-excess phenotype in the mesophyll and in the guard cells (Fulton et al., 2008; Horrer et al., 2016; Figure 1B-D). Thus, under the hypothesis of pool substitution between cell types, we would expect *bam3* to have similar transpiration dynamics as Col-0, and *bam1 bam3* to resemble *mex1* or *sex1*. However, both *bam3* and *bam1 bam3* behaved similarly as the wild type (Figure 8C-K, Supplemental Figures S13, S14, S15), and when marginal differences were detected they were not confirmed in experiment #3 (Supplemental Figures S4, S10). Hence, no direct connexion could be done between diel transpiration dynamics and starch turnover, either in the guard cells or in the rest of the leaf, suggesting more complex spatiotemporal effects.

It should be noted, though, that starch turnover and maltose production are not totally abolished in *bam1 bam3* (David et al., 2022), making it possible that a small fraction of sugars originating from mesophyll starch degradation diffuses into the apoplast, reaches the guard cells and sustains stomatal preopening as the night progresses. Based on gas exchange experiments in *Vicia faba*, it was suggested that a by-product of starch metabolism acts as a molecular link between daytime photosynthesis and stomatal preopening in the following night (Easlon and Richards, 2009). Could mobile maltose be such a signal? Consistent with this premise, maltose transport is detected in Arabidopsis leaves (Lu et al., 2006), while exogenous maltose added on epidermal peels is metabolized by guard cells (Dittrich and Raschke, 1977) and gently enhances stomatal opening (Dittrich and Mayer, 1978). However, feeding leaves with high concentration of exogenous maltose (1 mM) through the transpiration stream had no obvious short-term effect on stomatal dynamics, either in the dark, in the light, or during a dark-to-light transition (Supplemental Figure S16). An alternative candidate could be malate, which has been proposed to coordinate mesophyll photosynthesis and stomatal behaviour (Araújo et al., 2011; Daloso et al., 2017). However, two independent mutants impaired in ABCB14, a guard-cell malate importer (Lee et al., 2008) with normal starch turnover (Supplemental Figure S17), showed diel dynamics of transpiration that were virtually indistinguishable from that of the wild type (Figure 9, Supplemental Figure S18). Thus, we could not confidently conclude that either maltose or malate is a mobile signal that promotes nighttime stomatal preopening.

**Figure 9.**
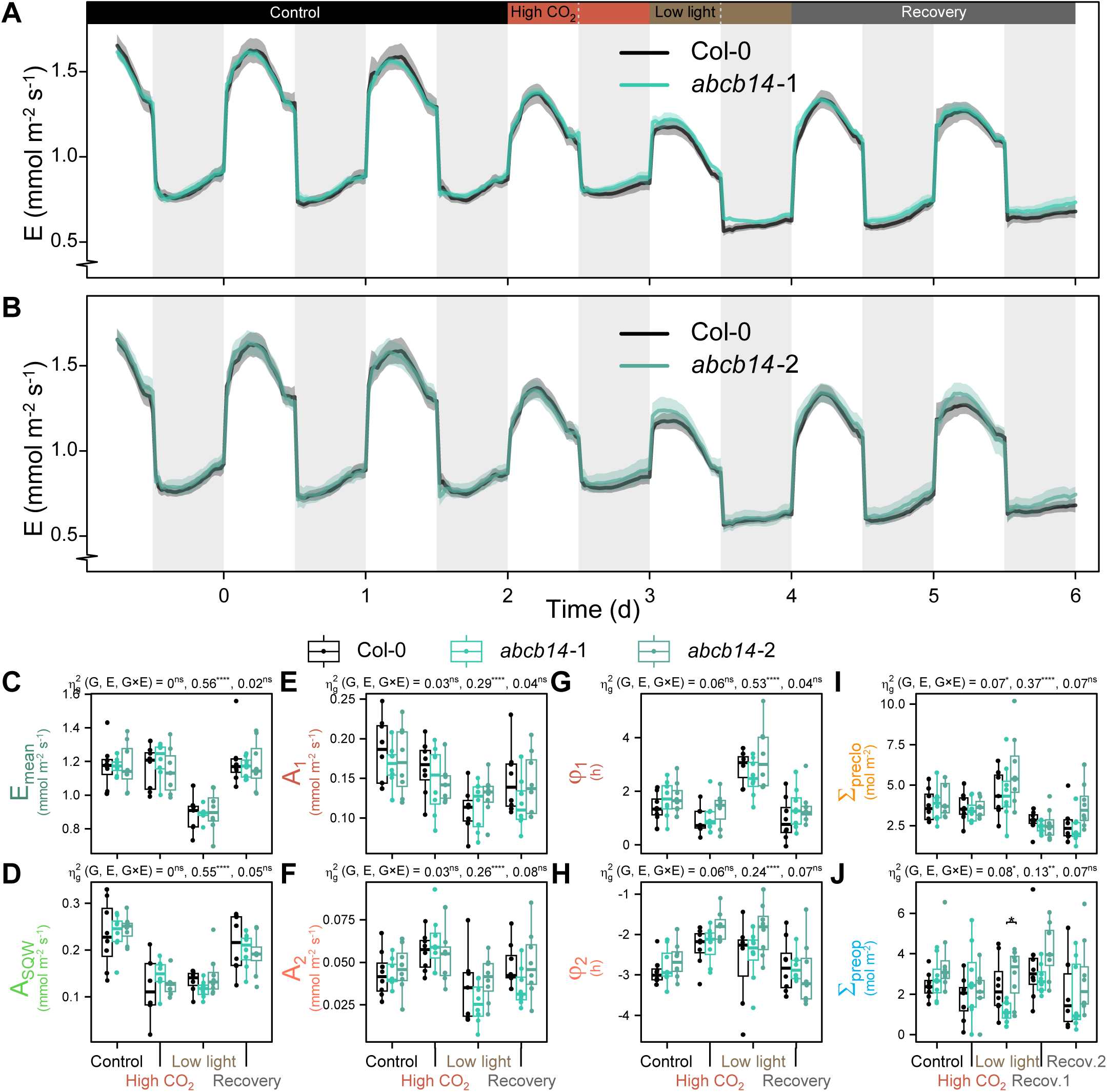
Analysis of diel transpiration of *abcb14-1 and abcb14-2*, two mutants impaired in a guard-cell malate importer. **(A)** and **(B)** Diel dynamics of transpiration rate of the wild type Col-0 compared to *abcb14-1* **(A)** and *abcb14-2* **(B)** in the same conditions as in Figure 3. The colour-shaded areas around the mean lines represent the means ± SE. **(C)** to **(H)** Boxplots of fitted parameters: **(C)** E_mean_, **(D)** A_SQW_, **(E)** A_1_, **(F)** A_2_, **(G)** φ_1_ and **(H)** φ_2_. **(I)** and **(J)** Boxplots of selected parameters extracted from the measured data: **(I)** Σ_preclo_ and **(J)** Σ_preop_. Abbreviations for parameters and significance codes for ANOVA effect sizes (Σ^2^_g_) and pairwise t-tests are like those in Figures 4 and 5.

## DISCUSSION

### The PhenoLeaks pipeline for monitoring the diel dynamics of transpiration highlights an endogenous stomatal rhythm with harmonic oscillations

Tailoring stomatal dynamics emerges as a realistic strategy for improving water use efficiency (Papanatsiou et al., 2019), but progress has been hindered by the lack of accessible phenotyping systems to monitor the genotypic variability for temporal patterns of transpiration. Here, we developed the PhenoLeaks pipeline that extends the capabilities of an existing phenotyping system initially designed to control soil water content by gravimetry (Granier et al., 2006), allowing us to monitor the diel transpiration dynamics on 150 pots with a 30-min resolution. We obtained similar dynamics as in previous Arabidopsis studies that used whole-plant gas exchange systems (Lascève et al., 1997; Sun et al., 2019), but also single leaf porometry (Matthews et al., 2018; Li et al., 2020) and direct measurements of stomatal aperture (Hassidim et al., 2017; Li et al., 2020). This indicates that the diel dynamics in transpiration are essentially driven by stomatal movements, while other factors potentially influencing the patterns of transpiration and also showing diel variations (Apelt et al., 2015), such as growth (that affects the leaf area by which transpiration is normalized) and hyponasty (that may affect the boundary layer conductance), are of secondary importance. Our calculations of transpiration assume constant growth rate of the transpiring area, not only because leaf expansion is not trivial to monitor at high temporal resolution, but also because we do not know how closely the acquisition of stomatal function paces tissue morphogenesis (*i*.*e*. the temporal mismatch between leaf area and transpiring area). Nonetheless, considering the available patterns of diel areal growth in Col-0 and *pgm* (Apelt et al., 2015) would make their diel dynamics to get closer together at night (after 4 h in the dark, growth of *pgm* is almost null while it only slowly decreases in Col-0), but to be still more delayed during the day (growth in Col-0 is temporarily dampened during the two first hours of the day, while that of *pgm* is boosted). Thus, delayed transpiration in *pgm* arises from stomatal behaviour. Overall, our setup was amenable for tracking diel stomatal rhythms of various genotypes over several days in contrasting environments. Differences between experiments were apparent, emphasizing the advantage of measuring many individuals simultaneously as made possible here. Moreover, across experiments, good reproducibility was obtained on the genotypic rankings for the measured and fitted parameters. The PhenoLeaks pipeline can readily be applied to any time-series of weight data obtained on any species.

PhenoLeaks also enabled us to provide a quantitative framework that accounts for the diel dynamics of transpiration, overarching daytime and nighttime stomatal movements and thereby hinting at the circadian clock. A previous model published by Matthews et al. (2018) and augmented by Vialet-Chabrand et al. (2021) was very powerful in accounting for both the rapid stomatal responses to light fluctuations and the endogenous daytime rhythm, but nighttime transpiration was not considered. Here, we propose a simple sinusoidal model that can be readily transformed and fitted through linear regression, with four key parameters (amplitudes A_1_ and A_2_, phases ϕ_1_ and ϕ_2_) to describe the endogenous rhythm of transpiration both day and night. Decomposition of a rhythmic signal into a series of periods, amplitudes and phases is a classical approach in the field of the circadian clock (Plautz et al., 1997). Studies in other organisms have revealed that the expression of many genes oscillates with a 24-h period but also with a 12-h period (Hughes et al., 2009), and the idea that the circadian clock may generate harmonic oscillations has gained theoretical and experimental support (Westermark and Herzel, 2013; Martins et al., 2016). In Arabidopsis like in many plants, there is strong diel rhythmicity for starch and sugar metabolism, and it is partly under circadian control (Bläsing et al., 2005; Lu et al., 2005). Moreover, the endogenous stomatal rhythm itself is under circadian control (Hubbard and Webb, 2015). Thus, that the rhythmic patterns of stomatal movements show harmonic oscillations makes biological sense. Nonetheless, the variations in ϕ_2_ and A_2_ were not clearly associated with a phenotype, and we cannot exclude that the second harmonic is just a convenient modelling solution that provides statistical flexibility to account for alternative mechanisms, whereby other actors (like ATP production, see below) actually gate the main circadian signal. Beyond the nature of the second harmonic, both observed and modelled data were consistent with the conclusion that mutations in starch metabolism affect the endogenous rhythm of stomata, likely by interfering with the circadian clock.

### Starch and sugars plausibly affect the endogenous rhythm of transpiration through interplay with the circadian clock

We propose that starch and sugars affect the timing of the endogenous rhythm of transpiration primarily through an interplay with the clock, while their energetic effect is usually non-limiting for stomatal movements as long as carbon starvation is precluded (Figure 10). Starch and sugar metabolism interacts with the clock in various ways. For instance, nighttime starch degradation is under circadian control (Graf et al., 2010). Conversely, in the dark, sucrose regulates the circadian clock through stabilization of its *GIGANTEA* component (Dalchau et al., 2011; Haydon et al., 2017). Moreover, in the early morning the clock is entrained by photosynthesis-derived sugars through the *PSEUDO-RESPONSE REGULATOR 7* component (Haydon et al., 2013). This is mediated by the transcription factor BASIC LEUCINE ZIPPER 63, itself regulated by the sugar-sensing kinase SnRK1 (Frank et al., 2018). Interestingly, sugars advance the phase of the clock in the morning, but delay it at night (Haydon et al., 2013), which would feedback on starch degradation rate and maintain carbon homeostasis throughout the diel cycle (Seki et al., 2017). Thus, abnormal sugar fluctuations triggered by mutations in the starch pathway (or by changes in the environment) may disturb the phase of the clock and the endogenous stomatal rhythm, with different effects depending on the timing and nature of the metabolic perturbations. For instance, functional interactions within the BAM family generate atypical patterns of maltose production in the *bam* mutants, so that *e*.*g*. at the start of the day, leaf maltose content is actually higher in *bam3* than in Col-0, while that in *bam1 bam3* is similar as in the wild type (David et al., 2022). This temporal complexity could subtend the variable transpiration phenotypes of the *bam* mutants if maltose, or a sugar derivative, shifts the endogenous stomatal rhythm by acting at a specific time of the diel cycle. Hence, subtle differences in the spatiotemporal patterns of sugar content, such as for *pgm vs*. *pgi*, or *sex1 vs*. *bam1 bam3*, may generate different phase shifts within the guard cells, underlying the inconsistent association between bulk-leaf starch turnover and stomatal movements.

**Figure 10.**
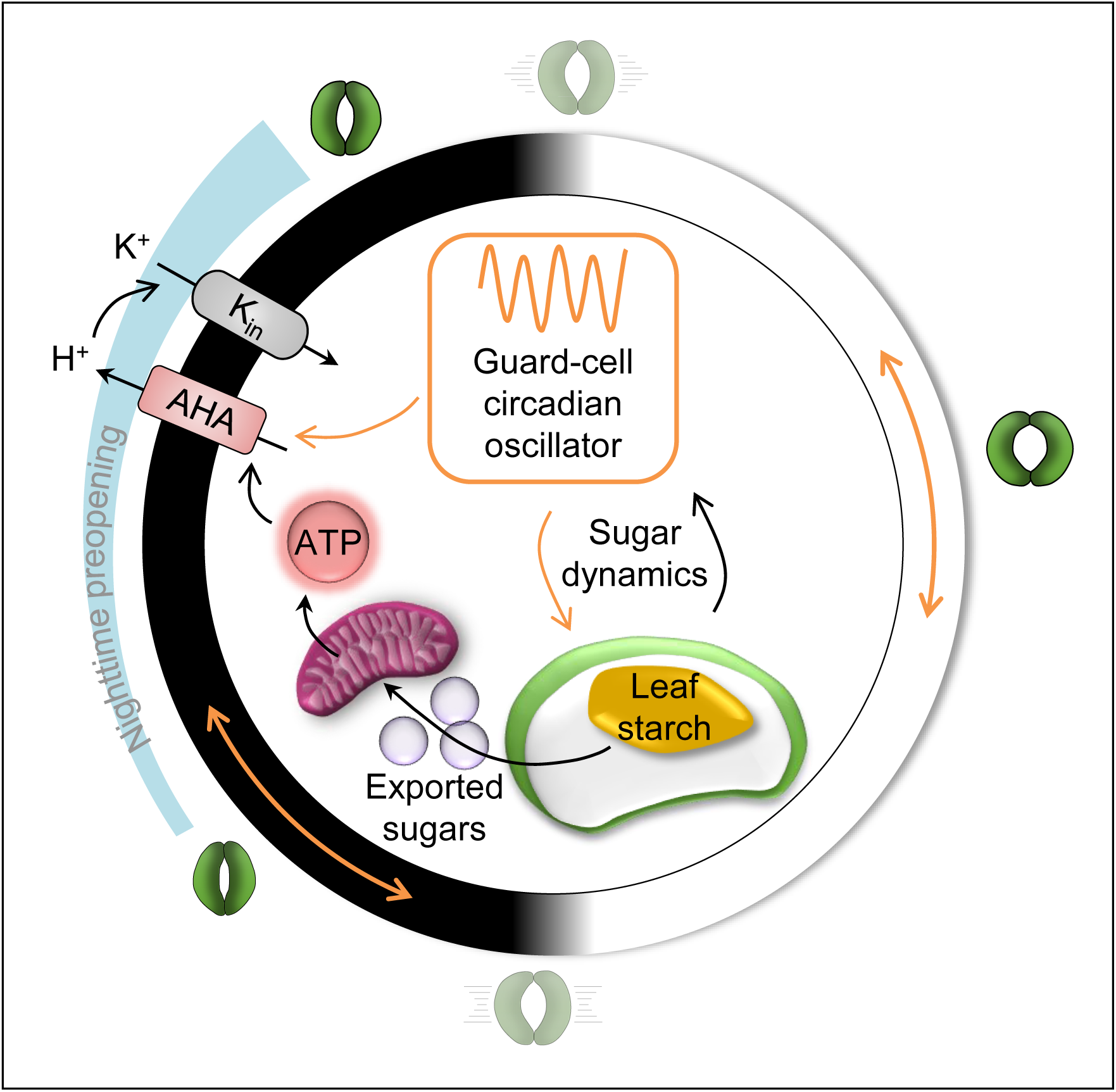
Conceptual model linking leaf starch metabolism and diel transpiration dynamics. Throughout the diel cycle (white and black circle), stomata show rapid movements at the day/night transitions, as well as an endogenous rhythm so that pore aperture reaches its maximum towards the middle of the day and its minimum in the early night. Starch metabolism in the leaf chloroplasts (either in the mesophyll or in the guard cells) influences sugar dynamics in the guard cells, which interacts with the circadian oscillator therein. The local clock then sets the time of maximal and minimal aperture of stomata (double-headed, curved orange arrows) by modulating the activity of the guard-cell H^+^-ATPase (AHA) that energizes ion transport (for instance K^+^ import through K_in_ inward channels) to drive stomatal opening. In turn, AHA requires energy in the form of ATP, which is usually provided by photosynthesis during the day and mitochondrial respiration of starch-derived sugars at night. In severe starch mutants showing starvation at night, stomatal preopening (blue funnel) is impaired due to the lack of respiratory ATP production. Note that the energetic effect of guard-cell starch at the night-to-day transition is not depicted.

How could an interplay between sugar metabolism and the clock ultimately modify stomatal movements? It is likely that sugars coming from the mesophyll add to the sugar pool of the guard cells, where they may adjust the local clock (that is tissue-specific; Endo, 2016) that gates ion transport together with ATP availability (Figure 10). An attractive candidate at the crossroads of sugar metabolism, circadian clock and ion transport in the guard cells is the florigen FT (FLOWERING LOCUS T), as well as its paralog TSF (TWIN SISTER OF FT). FT and TSF are normally involved in the photoperiodic control of flowering, and their induction requires trehalose-6-phosphate, a metabolic intermediate considered as a signal of carbon satiation (Wahl et al., 2013). In the guard cells, FT and TSF are controlled by several proteins involved in the circadian clock and photoperiodism (EARLY FLOWERING 3, GIGANTEA, CONSTANS) and gate the signalling pathway responsible for blue-light-induced stomatal opening (Kinoshita et al., 2011; Ando et al., 2013). Moreover, the *ft* mutant in continuous light has barely open stomata that are devoid of circadian rhythm (Kinoshita et al., 2011). The presence of FT, governed by the clock, would then permit light-activation of the plasma membrane H^+^-ATPase (Hubbard and Webb, 2011; Kinoshita et al., 2011). The activity of H^+^- ATPase energizes solute transport and impacts transpiration (Merlot et al., 2007; Blatt et al., 2014). H^+^-ATPase activity in the guard cells enhances the baseline of stomatal aperture in the daytime and the nighttime (Costa et al., 2015) and is a major limiting factor for light-induced stomatal opening (Wang et al., 2014), for instance by stimulating the activity of inward K^+^ channels, which are key effectors in stomatal opening (Jezek and Blatt, 2017). Interestingly, the *kincless* mutant, which is impaired in inward K^+^ current, displays defects in transpiration dynamics, with slowed daytime opening, impeded nighttime preopening, and impaired circadian rhythm (Lebaudy et al., 2008). Thus, in starch mutants such as *pgm*, *isa1*, *sex1* or *mex1*, strong metabolic disorders – including in the mesophyll – that eventually upset sugar content within the guard cells may locally translate into a phase shift in the circadian clock, which would then delay FT production, H^+^-ATPase activity, K^+^ import, turgor build-up and stomatal (pre)opening. In turn, the activity of the proton pump requires energy in the form of ATP, which may originate from starch degradation products and is thus likely to be limiting in the most severe starch mutants at night.

### The energetic role of starch degradation products is usually not a limiting factor for the endogenous stomatal rhythm, but is emergent upon severe starvation

While no instant relationship was observed between sugar content and stomatal movements in the daytime, the mutations inducing lowest sugar availability at night broadly coincided with an impaired nighttime preopening. These mutations included *pgm* that cannot synthesize starch, *isa1* that stores labile phytoglycogen instead of starch, *sex1* that cannot initiate nighttime starch breakdown, and *mex1* that cannot transfer maltose outside the chloroplast. The fact that stomatal preopening may be observed in situations where leaf starch turnover is severely reduced, such as in the *pgi* and *bam1 bam3* mutants or under low light, could then be explained by a residual starch turnover being sufficient to fuel ATP production in the guard cells. Thus, severe starvation is required to observe the ATP-limitation over stomatal movements.

The idea that carbon starvation precludes stomatal preopening at night is in line with the observation that mitochondrial respiration releases the main source of energy for stomatal movements when photosynthesis is null or low (Vialet-Chabrand et al., 2021). Notably, both light-induced stomatal opening and darkness-induced stomatal closure are potentiated by O_2_ availability, highlighting a key role for respiratory processes in the control of stomatal movements (Vialet-Chabrand et al., 2021). We suggest that nighttime endogenous preopening is also limited by respiration, which generates ATP from starch degradation products. In our model, the circadian clock poises the potential activity of H^+^-ATPase, which is then empowered by actual ATP availability (Figure 10). This dual control can be illustrated in *pgm*, where respiration is tightly correlated with sugar content (Rasse and Tocquin, 2006), being high during the day and at the onset of the night, after which respiration decreases rapidly for the rest of the night – when that of Col-0 is virtually constant (Caspar et al., 1985; Lascève et al., 1997; Pantin, 2011). Thus, in *pgm*, the shifted clock delays stomatal closure in the early night, while the lack of respiratory ATP production in the absence of starch impedes preopening as the night progresses.

Starvation could potentially prevent stomatal movements across the whole diel cycle, as observed in seedlings grown in plates, where very low carbon availability inactivates TARGET of RAPAMYCIN (TOR, a kinase involved in energy signalling), which leads to BAM1 inactivation, abolishment of starch degradation in the guard cells, and loss of endogenous stomatal rhythm (Han et al., 2022). In our study, neither *bam1* nor *amy3 bam1* had a discernible transpiration phenotype, suggesting cell-autonomous starch breakdown is dispensable to maintain sugar homeostasis and endogenous rhythm in the guard cells of more mature, less starved plants. Beyond their effects on the energy availability or the circadian clock, fluctuations in sugar content may also indirectly disrupt the endogenous rhythm of transpiration. For instance, transient periods of carbon starvation may ‘imprint’ the metabolism or ion transport system of the guard cells, so that stomata are not able to respond to sugars immediately after resumption. Supporting this idea, critically low levels of sugars have strong effects on the diel transcriptome (Gibon et al., 2004b; Bläsing et al., 2005), and every night in *pgm* such starvation impacts gene expression and enzyme activities in the following day (Gibon et al., 2004a, 2004b). On the other hand, there could be a temporal mismatch between the sugar contents measured at the bulk leaf level and the amount of sugars actually available for metabolism within the guard cells. For instance, the expression of *STP1* is enhanced at night but repressed by sugars (Stadler et al., 2003; Cordoba et al., 2015), making it possible that the sugar import system of the guard cells is out of phase with apoplastic sugar availability in the most severe starch mutants. Overall, spatiotemporal uncoupling between sugar metabolism and stomatal movements may arise from different mechanisms, making it hard to predict the transpiration phenotype of a given mutant based on its starch turnover or sugar content in the absence of a quantitative, mechanistic model (Blatt et al., 2022).

## CONCLUSION

Several starch mutants showed a phase shift in the endogenous stomatal rhythm, and acute starvation at night coincided with impaired nighttime preopening. However, neither tissue-specific starch turnover nor instant sugar content were reliable predictors of stomatal movements at any time of the diel cycle. In our conceptual framework (Figure 10), leaf starch metabolism affects sugar availability in the guard cells, which modulates the local circadian clock. In turn, the clock sets the potential activity of the guard-cell H^+^-ATPase that empowers guard-cell ion transport, turgor build-up and diel stomatal movements. At night, minimal starch breakdown fuels the production of respiratory ATP that is basically required to empower this process. Our model highlights a dual role for starch-derived sugars in setting the diel tempo of the guard cells and energizing ion transport therein.

## METHODS

### Plant material

Leaf starch metabolism in Arabidopsis is largely documented (Streb and Zeeman, 2012). All starch mutants used in this study were described previously and in the Col-0 background: *pgm*-1 (AT5G51820; Caspar et al., 1985), *pgi*-1 (AT4G24620; Yu et al., 2000), *isa1*-1 (AT2G39930; Delatte et al., 2005), *ss4*-3 (AT4G18240; Crumpton-Taylor et al., 2013), *sex1*-3 (AT1G10760; Caspar et al., 1991; Yu et al., 2001; Ritte et al., 2002), *amy3*-2 (AT1G69830; Yu et al., 2005), *bam1* (AT3G23920; Fulton et al., 2008), *bam3* (AT4G17090; Fulton et al., 2008), *amy3*-2 *bam1* (Horrer et al., 2016; Flütsch et al., 2020b), *bam1 bam3* (Fulton et al., 2008; Horrer et al., 2016; David et al., 2022), *mex1*-1 (AT5G17520; Niittylä et al., 2004), *dpe1*-2 (AT5G64860; Critchley et al., 2001) and *dpe2*-5 (AT2G40840; Chia et al., 2004). For sake of clarity, the allele number of the starch mutants has been omitted throughout the text and figures. Due to seed availability, the lines of *isa1*-1 and *ss4*-3 we used also contained a starvation reporter construct consisting of the promoter of the starvation gene AT1G10070 fused to luciferase coding sequence (*ProStarv::LUC*; Graf et al., 2010) that was inserted in the original mutants as well as in Col-0, as described by Crumpton-Taylor (2010). The transformed lines were denoted with a dagger throughout the figures. No difference in the transpiration pattern was observed between Col-0 and Col-0^†^. We also used two mutants affected in a guard cell malate transporter, *abcb14*-1 and *abcb14*-2 (AT1G28010; Lee et al., 2008).

### Growth conditions

Four independent experiments were performed: #1 and #2 comprised only Col-0, while #3 was a pilot experiment that also included some starch mutants (*pgm*, *sex1*, *bam3*, *bam1 bam3*) before the main experiment #4, which included all genotypes (Supplemental Table S1). The plants were grown and assayed in one of the three Phenopsis phenotyping platforms (Granier et al., 2006). Arabidopsis seeds were sown in 200-ml cylindrical plastic pots filled with a weighed amount of soil, and then thinned to one plant per pot. Eight plants of each genotype were obtained. To analyse all genotypes on the same day while reducing differences in plant architecture arising from contrasting growth rates among genotypes, the sowing date was adapted for each genotype based on our previous experience with starch genotypes (Pantin et al., 2013a) for the pilot experiment #3, and then adjusted for the main experiment #4 (Supplemental Table S1). Plants were then grown under 12-h photoperiod at an irradiance of 180 µmol m^-2^ s^-1^ and ambient atmospheric CO_2_ (∼400 ppm). In experiment #4, air temperature (22°C), relative humidity (66%) and vapour pressure deficit (0.9 kPa) were constant during day and night, and eight plants per genotype were obtained (see Supplemental Table S1 for the other experiments). Each pot was weighed daily and its soil water content kept constant (1.4 g_water_ g^-1^) by watering with a modified Hoagland solution. In experiment #4, a moderate soil water deficit was also applied on eight independent plants for Col-0 and *pgm*. Irrigation was withheld 10 d before the start of the transpiration assay, and the target soil water content of 0.75 g_water_ g^-1^ was reached within 4 d.

### The PhenoLeaks pipeline to monitor transpiration dynamics

#### Gravimetry experiment

Transpiration was measured by gravimetry. For the assay, the surface of each pot was covered with cellophane to prevent evaporation from the soil, and then with a plastic grid to facilitate further image analysis. After a short stabilization period, each pot was weighed at a frequency of about 30 min by a moving scale (0.001 g precision, Precisa XB 620M, Precisa Instruments AG, Dietikon, Switzerland) for a duration that depended on the experiment. When necessary, irrigation was performed manually using a syringe in order to maintain the target soil water content individually for each pot. As explained in the results, we noticed *a posteriori* that the largest plants from experiment #3 showed deviation from the target (Figure 2, Supplemental Figure S2). For experiment #4, after the control period (in the same environmental conditions as for growth) that lasted for about 3 d, atmospheric CO_2_ was doubled to 800 ppm during the daytime, and returned to 400 ppm at night. At the start of the next day, irradiance was reduced to 50 µmol m^-2^ s^-1^ during the daytime. Then plants were given 2 d of recovery in the control, growth conditions. Environmental data were recorded together with the pot weights in the PhenopsisDB database (Fabre et al., 2011).

#### Data processing

The gravimetrical data were then processed using R scripts. Manual irrigation events and other outliers were detected by analysing the studentized residuals of a local linear regression model of pot weight loss against time. Afterwards, the regression was run again and individual pot transpiration was calculated at every 30 min (falling on the day/night transitions) as the fitted slope of the local linear relationship. Because moving linear regression inherently buffers rapid changes such as occurring during the day/night transitions, we made sure that the time interval used for the local regression did not span these transitions. This was achieved by setting a variable interval depending on the current position in the day/night cycle: at the start of each day or night period, the interval was set to 90 min (∼3 points), gradually increased to 360 min (∼12 points) and then gradually decreased to 120 min (∼4 points) at the end of the same period. This varying interval allowed us to achieve a satisfactory trade-off between temporal resolution at the day/night transitions and robustness elsewhere. The rate of water loss was corrected for plant growth to obtain transpiration per unit of projected leaf area. Leaf area was estimated by a linear or polynomial model (automatically selected for each plant depending on the fit quality) based on measurements of the daily projected surface extracted by image analysis. A script was developed to detect punctual outliers that may occasionally be generated by mechanical instability of the balance. If the issue persisted for several hours in a given pot, the whole diel cycle involved was discarded to avoid bias in estimating the parameters. Note that the actual number of replicates available for each genotype and for each period can be visualized as the number of points in each corresponding jitter plot.

#### Extraction of parameters

To quantify the dynamics, we extracted one set of parameters per plant and per environmental condition. The data were first linearly detrended by using the control conditions (and recovery for experiment #4) to estimate the slope due to ageing (calculated of the mean of the slope of the linear regressions of transpiration against time performed separately for the daytime and the nighttime). For the control conditions, we first averaged the initial 2.75 d into a single 24-h kinetics. We computed the mean transpiration over 24-h (E_diel_) and over each period (E_day_, E_night_), as well as dynamic parameters (Figure 4A). We estimated the diel amplitude of transpiration (A_diel_), the time of maximal daytime transpiration (t_day max_), the cumulative afternoon preclosure after t_day max_ (Σ_preclo_), the time of minimal nighttime transpiration (t_night min_) and the cumulative nighttime preopening after t_night min_ (Σ_preop_). Then, we separated the rapid changes observed at the day/night transitions from the endogenous changes of the rest of each period. The exponential response of stomata to a rapid change in light intensity have largely plateaued after 30 min (Matthews et al., 2018). Even in “delayed genotypes” such as *pgm* and *mex1*, the changes observed after 30 min were not quantitatively consistent with a prolongation of the exponential kinetics. Therefore, the magnitude of rapid opening at the night-to-day-transition (A_rapid op_) and that of the rapid closure at the day-to-night transition (A_rapid clo_) were estimated at 30 min after a transition. Finally, we estimated the average slope of change in transpiration throughout the daytime after 30-min illumination (σ_day_), the cumulative transpiration dynamics above this daytime trend during the same period (Δ_day_, usually positive; the area below the trend, if any, being subtracted), the average slope of change in transpiration throughout the nighttime after 30-min darkness (σ_night_) and the cumulative transpiration dynamics below this nighttime trend during the same period (Δ_night_, usually negative; the area above the trend, if any, being added). The same procedure was applied on the fitted curves, and the obtained parameters were denoted with a hat (Figure 5B). Cumulative parameters were obtained through trapezoidal integration for the measured data (Σ_preclo_, Σ_preop_, Δ_day_,

Δ_night_), and by calculating primitive integrals for the fitted curves (Σ_preclo_, Σ_preop_, Δ_day_, Δ_night_). The code will be made available as R scripts on the PhenoLeaks repository (GitHub). The link will be provided upon publication of the manuscript in a peer-reviewed journal.

### Modelling the diel transpiration dynamics

#### Model structure

To model the transpiration dynamics, we used a periodic signal (Figure 5A) composed of a square wave (period fixed to 24 h, semi-amplitude A_SQW_, phase ε), a fundamental sine wave (period fixed to 24 h, semi-amplitude A_1_, phase ϕ_1_) and its second harmonic (period fixed to 12 h, semi-amplitude A_2_, phase ϕ_2_) that oscillates around a baseline transpiration (E_mean_):

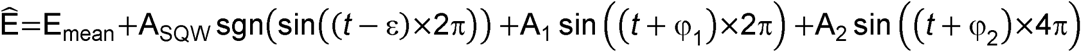

with *t* the time (*t* = 0 being the start of the day period of the first complete diel cycle) and sgn() the signum function. We shifted the phase of the square wave by ε = −15 min (*i*.*e*. the square wave lags 15 min behind the day/night transitions) so that the square wave is +A_SQW_ during the day and at the day-to-night transition, −A_SQW_ during the night and at the night-to-day transition, and 0 at 15 min after the transitions (when there is no data to be fitted since transpiration is estimated at every 30 min starting from 0).

In the model augmented for acclimation, we implemented a linear pulse that varies linearly over 24 h within a change in the environmental conditions:

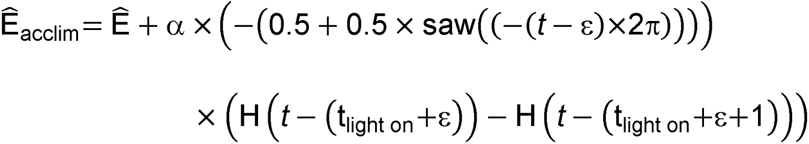

with H() the Heaviside step function, saw() the sawtooth function, t_light on_ the time (d) when the light switches on in the given acclimation period, and α the acclimation slope, so that the acclimation pulse is –α at the start of the day when the environment changes, and 0 at the end of the following night, ensuring continuity if the new environment extends beyond one diel cycle (*i*.*e*. recovery in experiment #4 and low light in experiment #3).

#### Model fitting

To fit the first model to the data, we transformed the model so that it can be fitted using multiple linear regression:

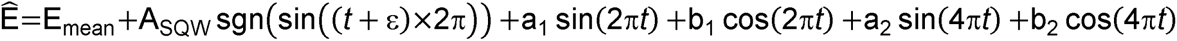

with:

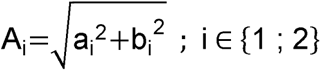

and

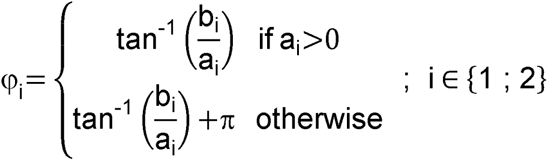

Once the parameters were obtained, the phases were wrapped so that −12 h < ϕ_1_ ≤ +12 h and −6 h < ϕ_2_ ≤ +6 h.

For each plant, a set of parameters was initially obtained using data from the interval of each period of interest. In experiment #4, we used] −0.5 d; 2 d] for the control conditions,] 2 d; 3 d] for elevated CO_2_,] 3 d; 4 d] for low irradiance,] 4 d; 5 d] for recovery cycle one, and] 5 d; 6 d] for recovery cycle two. In experiment #3, we used] −0.5 d; 1 d] for the control conditions (avoiding excessive soil drying), and] 3 d; 4 d] for low irradiance (with the fast soil drying pots identified in Supplemental Figure S2A being discarded from the analysis).

The same procedure was implemented for the acclimation model, just adding the acclimation term to fit α. In experiment #4, both recovery days were fitted jointly, so that they shared the same set of parameters but only the first recovery cycle was affected by the 24-h linear pulse. Similarly, in experiment #3, the data used to fit the low light parameters extended to the second day of low light.

The code will be made available as R scripts on the PhenoLeaks repository (GitHub). The link will be provided upon publication of the manuscript in a peer-reviewed journal.

### Gas exchange for maltose feeding assays

Gas exchange experiments were performed on xylem-fed detached leaves as described previously (Pantin et al., 2013b). Mature leaves of Col-0 plants were excised one hour before the end of the night period (darkness experiment) or during the daytime (light experiment), and immediately recut under water. The petiole was then dipped in an Eppendorf containing water and the leaf was inserted into a 2×3 LED chamber of an Li6400-XT infrared gas analyser (Li-Cor Inc., Lincoln, NE, USA) connected to an Li610 dew-point generator (Li-Cor Inc.), and data were logged every 10 s. CO_2_ was set at 400 ppm, block temperature at 22°C and reference dew-point temperature at 17°C. Irradiance was set at either 0, 50, 200 or 500 µmol m^-2^ s^-1^ depending on the experiment. Once stomatal conductance had stabilized, water was changed against a 1 mM maltose solution. For the maltose effect at the dark-to-light transition, leaves were harvested at the end of the night, dipped into 1 mM maltose or water (control) for at least 2 h in darkness before monitoring gas exchange at the dark-to-light transition.

### Starch analyses

During experiment #4, leaf samples for starch analysis were obtained at the end of the day and night periods on distinct plants. Four independent biological replicates were harvested per genotype at each time point. Approximately 40 mg of fresh leaf material per rosette was harvested, weighed and immediately frozen in liquid nitrogen. Starch content was analysed by enzymatic assay using spectrophotometric analysis of the residual fraction of ethanol-water extracts (Hummel et al., 2010).

For iodine staining of starch, leaves were harvested from the same batch of plants and decolourised in 80% (v/v) ethanol. Excess of ethanol was removed by rinsing with water. Leaves were stained with a 30% (v/v) lugol solution for 30 min followed by a wash step of 30 min with water. The iodine staining was observed using an Olympus BX61 (60x, guard cells) and SZX16 (0.5x, whole leaves) microscope.

### Statistical analyses

Data management, statistical analyses and figures were produced using R 4.2.0 (R Development Core Team, 2022). Starch contents showed strong heteroscedasticity and were analysed using the Kruskal-Wallis non-parametric test at the 5% level followed by multiple comparisons of the ranks through Fisher’s LSD adjusted by the Benjamini-Hochberg method (de Mendiburu, 2021). The parameters extracted from the transpiration dynamics were analysed using one-, two- or three-way ANOVA (note that mixed ANOVA was not used to avoid listwise deletion, *i.e.* the complete removal of one pot if only one period is missing). ANOVA effect sizes of individual factors and their interactions were computed using the generalized eta squared (η^2^), which was followed by pairwise t-tests adjusted by the Bonferroni method (Kassambara, 2021). The figures systematically show the significant differences between genotypes for each environment. When required, similar tests were carried out to evaluate the significance of the differences between environments, which was reported in the main text.

### Accession numbers

Sequence data from this article can be found in the EMBL/GenBank data libraries under accession numbers AT5G51820 (*PGM*), AT4G24620 (*PGI*), AT1G10760 (*SEX1*/*GWD*), AT1G69830 (*AMY3*), AT3G23920 (*BAM1*), AT4G17090 (*BAM3*), AT5G17520 (*MEX1*), AT5G64860 (*DPE1*), AT2G40840 (*DPE2*), AT4G18240 (*SS4*), AT2G39930 (*ISA1*) and AT1G28010 (*ABCB14*).

## Supplemental data

**Supplemental Table S1.** Main conditions in the four experiments.

**Supplemental Figure S1.** Statistical comparison of transpiration dynamics on experiments #1 to #4 for Col-0 in control conditions.

**Supplemental Figure S2.** Characterization of the plants with varying soil water content in experiments #3 and #4.

**Supplemental Figure S3.** Comparison of further transpiration parameters for *pgm* and *sex1*.

**Supplemental Figure S4.** Analysis of diel transpiration of several starch mutants from the pilot experiment #3 and statistical comparison with the main experiment #4.

**Supplemental Figure S5.** Examples of individual fits in Col-0.

**Supplemental Figure S6.** Examples of individual fits in *pgm*.

**Supplemental Figure S7.** Examples of individual fits in *sex1*.

**Supplemental Figure S8.** Effect of the environment and the genotype on the fitted acclimation slope.

**Supplemental Figure S9.** Effect of the irrigation regime, the environment and the genotype on the fitted parameters.

**Supplemental Figure S10.** Effect of the experiment, the environment and the genotype on the fitted parameters.

**Supplemental Figure S11.** Comparison of further transpiration parameters for *isa1* and *ss4*.

**Supplemental Figure S12.** Comparison of further transpiration parameters for *mex1*, *dpe1* and *dpe2*.

**Supplemental Figure S13.** Comparison of further transpiration parameters for *pgi*, *amy3 bam1* and *bam1 bam3*.

**Supplemental Figure S14.** Analysis of diel transpiration of *amy3*, *bam1* and *bam3* single mutants.

**Supplemental Figure S15.** Comparison of further transpiration parameters for *amy3*, *bam1* and *bam3*.

**Supplemental Figure S16.** Effect of exogenous maltose on stomatal conductance of Col-0.

**Supplemental Figure S17.** Characterization of the starch patterns in *abcb14*-1 and *abcb14*-2.

**Supplemental Figure S18.** Comparison of further transpiration parameters for *abcb14*-1 and *abcb14*-2.

## ACKNOWLEDGEMENTS

We thank Jérémy Cortello and Alexis Bédiée for technical help during the transpiration experiments, Gaëlle Rolland for assistance during starch measurements, Carine Alcon for guidance in microscopy, Anna Medici for advices on genotyping, and the groups of Samuel C. Zeeman and Youngsook Lee for providing seeds of starch and *abcb14* mutants, respectively. We are grateful to Alison M. Smith and Doreen Feike for the gift of the LUC-expressing lines and advice on their analysis. We also acknowledge two anonymous reviewers for proposing new insight in the data interpretation and helping us to improve the manuscript. This research was supported by INRAE and Région Occitanie (PhD fellowship to A.J.W.).

## AUTHOR CONTRIBUTIONS

FP, TS and AJW designed the research; AJW and MD performed the transpiration experiments; FP performed the gas exchange experiments; AJW and FP developed PhenoLeaks; FP developed the model; AJW and FP analyzed data; FP wrote the draft of the paper; AJW and TS revised the manuscript.

