## Supplemental Data for "Leaf starch metabolism sets the phase of stomatal rhythm"

**Supplemental Table S1.** Main conditions in the four experiments.

**Supplemental Figure S1.** Statistical comparison of transpiration dynamics on experiments #1 to #4 for Col-0 in control conditions.

**Supplemental Figure S2.** Characterization of the plants with varying soil water content in experiments #3 and #4.

**Supplemental Figure S3.** Comparison of further transpiration parameters for *pgm* and *sex1*.

**Supplemental Figure S4.** Analysis of diel transpiration of several starch mutants from the pilot experiment #3 and statistical comparison with the main experiment #4.

**Supplemental Figure S5.** Examples of individual fits in Col-0.

**Supplemental Figure S6.** Examples of individual fits in *pgm*.

**Supplemental Figure S7.** Examples of individual fits in *sex1*.

**Supplemental Figure S8.** Effect of the environment and the genotype on the fitted acclimation slope.

**Supplemental Figure S9.** Effect of the irrigation regime, the environment and the genotype on the fitted parameters.

**Supplemental Figure S10.** Effect of the experiment, the environment and the genotype on the fitted parameters.

**Supplemental Figure S11.** Comparison of further transpiration parameters for *isa1* and *ss4*.

**Supplemental Figure S12.** Comparison of further transpiration parameters for *mex1*, *dpe1* and *dpe2*.

**Supplemental Figure S13.** Comparison of further transpiration parameters for *pgi*, *amy3 bam1* and *bam1 bam3*.

**Supplemental Figure S14.** Analysis of diel transpiration of *amy3*, *bam1* and *bam3* single mutants.

**Supplemental Figure S15.** Comparison of further transpiration parameters for *amy3*, *bam1* and *bam3*.

**Supplemental Figure S16.** Effect of exogenous maltose on stomatal conductance of Col-0.

**Supplemental Figure S17.** Characterization of the starch patterns in *abcb14*-1 and *abcb14*-2.

**Supplemental Figure S18.** Comparison of further transpiration parameters for *abcb14*-1 and *abcb14*-2.

**Supplemental Table S1.** Main conditions in the four experiments. Temp.: air temperature; RH: air relative humidity; VPD: air vapour pressure deficit; DAS: days after sowing. The sample size for Col-0 and *pgm* in experiment #4 is twofold due to the additional condition of stable soil water deficit. Supports Figure 2 and Supplemental Figures S1, S2, S4, S10.

| Experiment # (PhenopsisDB) | Growth chamber ID | Assay chamber ID | Temp. (°C) | RH (%) | VPD (kPa) | Genotype | Age at t0 (DAS) | Number of plants |
|---|---|---|---|---|---|---|---|---|
| #1 (C3M31) | Phenopsis3 | Phenopsis3 | 20/17 (day/night) | 68/62 (day/night) | 0.75 | Col-0 | 41 | 9 |
| #2 (C2M43A) | Phenopsis2 | Phenopsis2 | 22 | 62 | 1 | Col-0 | 30 | 6 |
| #3 (C2M43B) | Phenopsis2 | Phenopsis1 | 22 | 66 | 0.9 | Col-0 | 42 | 7 |
| | | | | | | *pgm* | 52 | 7 |
| | | | | | | *sex1* | 52 | 4 |
| | | | | | | *bam3* | 45 | 4 |
| | | | | | | *bam1 bam3* | 45 | 5 |
| #4 (C2M47) | Phenopsis2 | Phenopsis2 | 22 | 66 | 0.9 | Col-0 | 29 | 8 × 2 |
| | | | | | | Col-0† | 29 | 8 |
| | | | | | | *pgi* | 36 | 8 |
| | | | | | | *pgm* | 43 | 8 × 2 |
| | | | | | | *isa1* † | 29 | 8 |
| | | | | | | *ss4* † | 36 | 8 |
| | | | | | | *sex1* | 43 | 8 |
| | | | | | | *amy3* | 29 | 8 |
| | | | | | | *amy3 bam1* | 29 | 8 |
| | | | | | | *bam1* | 29 | 8 |
| | | | | | | *bam3* | 36 | 8 |
| | | | | | | *bam1 bam3* | 36 | 8 |
| | | | | | | *mex1* | 43 | 8 |
| | | | | | | *dpe1* | 36 | 8 |
| | | | | | | *dpe2* | 36 | 8 |
| | | | | | | *abcb14* -1 | 29 | 8 |
| | | | | | | *abcb14* -2 | 29 | 8 |

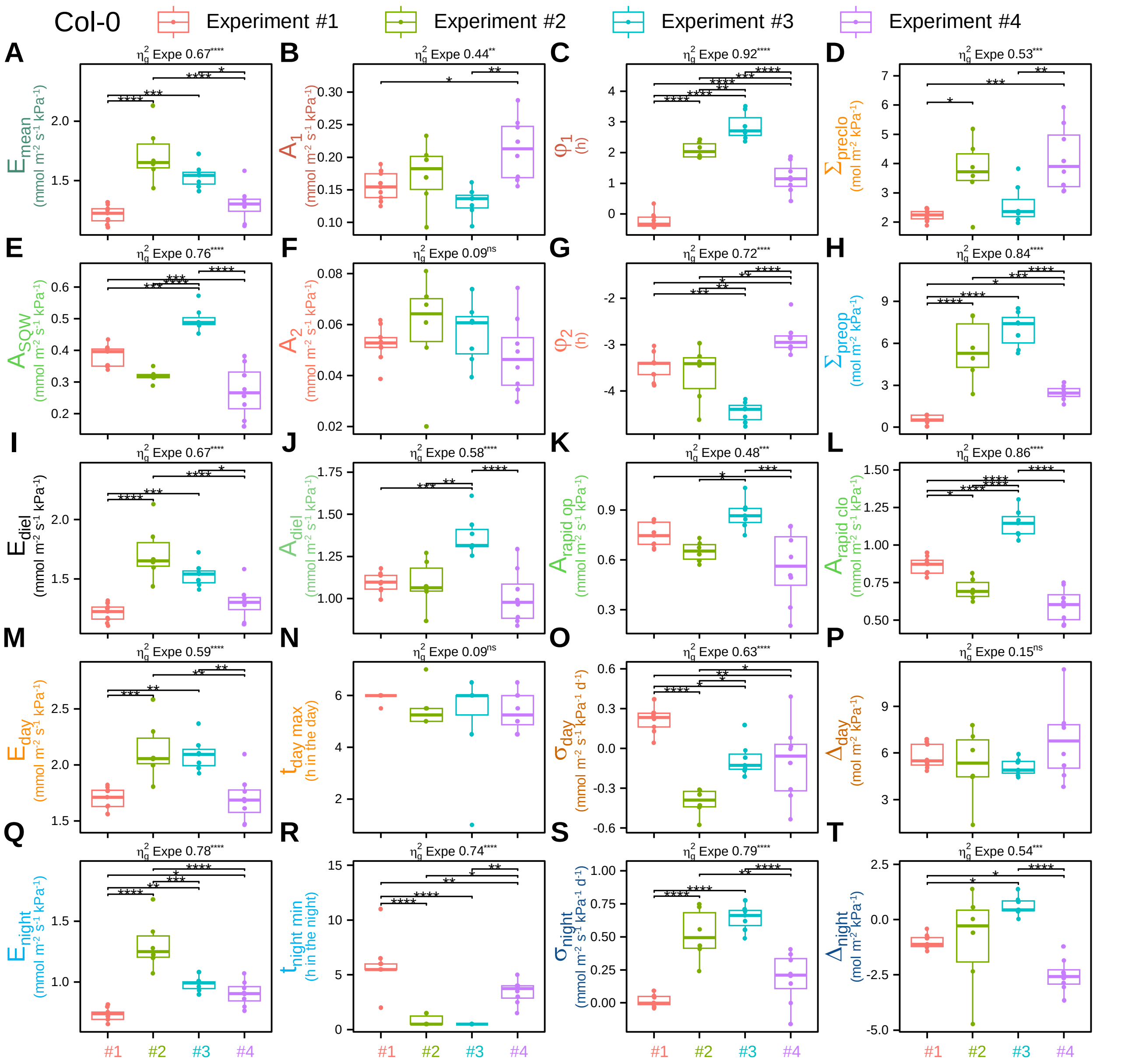


**Supplemental Figure S1.** Statistical comparison of transpiration dynamics on experiments #1 to #4 for Col-0 in control conditions. Different parameters were either fitted (A, B, C, E, F, G) or observed (the others) from the dynamics as explained in Figures 4A and 5A. **(A)** to **(T)** Boxplots of: **(A)** Mean transpiration rate over a 24-h period ($E_{mean}$), **(B)** semi-amplitude of the fundamental sine wave ($A_1$), **(C)** phase of the fundamental sine wave ($\varphi_1$), **(D)** cumulative afternoon preclosure after $t_{day\ max}$ ($\Sigma_{preclo}$), **(E)** semi-amplitude of the square wave ($A_{SQW}$), **(F)** semi-amplitude of the second harmonic ($A_2$), **(G)** phase of the second harmonic ($\varphi_2$), **(H)** cumulative nighttime preopening after $t_{night\ min}$ ($\Sigma_{preop}$), **(I)** mean transpiration over 24 h ($E_{diel}$), **(J)** diel amplitude of transpiration ($A_{diel}$), **(K)** rapid stomatal opening at the day-to-night transition ($A_{rapid\ op}$), **(L)** rapid stomatal closure at the night-to-day transition ($A_{rapid\ clo}$), **(M)** mean transpiration over the daytime ($E_{day}$), **(N)** time of maximal daytime transpiration ($t_{day\ max}$), **(O)** average slope of change in transpiration throughout the daytime after 30-min illumination ($\sigma_{day}$), **(P)** cumulative transpiration dynamics above the daytime trend ($\Delta_{day}$), **(Q)** mean transpiration over the nightime ($E_{night}$), **(R)** time of minimal nighttime transpiration ($t_{night\ min}$), **(S)** average slope of change in transpiration throughout the nighttime after 30-min darkness ($\sigma_{night}$), and **(T)** cumulative transpiration dynamics below the nighttime trend ($\Delta_{night}$). For each parameter, a one-way ANOVA was done and the effect size ($\eta^2_g$) of the experiment (Expe), as well as its significance level, are reported above each corresponding plot. Pairwise t-tests were systematically done to assess the differences between experiments, and the differences are reported on the plot when significant. Significance codes are as follows: $^{ns}$, not significant; *, $0.05 \leq p_{val} < 10^{-2}$; **, $10^{-2} \leq p_{val} < 10^{-3}$; ***, $10^{-3} \leq p_{val} < 10^{-4}$; ****, $10^{-4} \leq p_{val}$. Supports Figure 1.

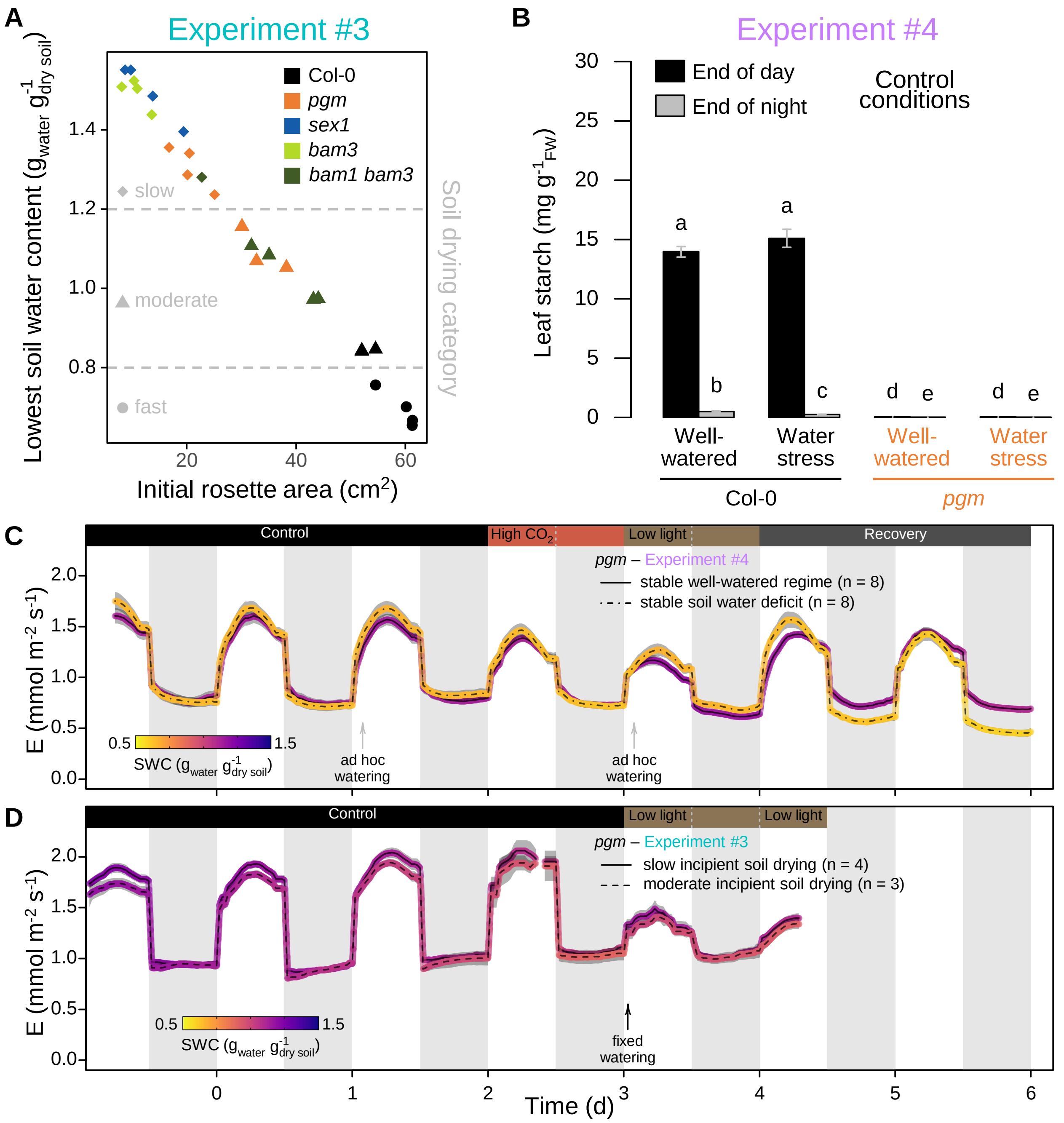


**Supplemental Figure S2.** Characterization of the plants with varying soil water content in experiments #3 and #4. **(A)** Relationship between rosette area at the start of the transpiration assay and the minimal soil water content (SWC) observed for each pot in experiment #3, where the target irrigation could not be adequately maintained. Based on the lowest SWC measured during the assay, each plant was *a posteriori* assigned one category (slow, moderate or fast soil drying). It was mainly driven by leaf area and thus unbalanced between genotypes. **(B)** Leaf starch content determined at the end of the day and night periods in control conditions of experiment #4 for Col-0 and *pgm* under well-watered conditions or stable soil water deficit. Error bars are means ± SE. Letters denote significant differences after a Kruskal-Wallis test (α = 5%) followed by multiple comparisons of ranks. The data for well-watered conditions are the same as the ones used in Figure 1B. **(C)** and **(D)** Transpiration dynamics of *pgm* throughout consecutive diel cycles in varying environmental conditions, for experiment #4 with two different stable watering regimes **(C)** and for experiment #3 with fixed watering irrespective of plant size **(D)**. Transpiration is colour-coded for the average SWC of each condition. The environmental conditions during the assay were the same as in Figure 2C and 2D. Note that SWC in experiment #3 was much less affected by fixed watering than in Col-0 (Figure 2D) due to smaller rosette area (panel A). The shaded areas around the mean lines represent the means ± SE. Supports Figure 2.

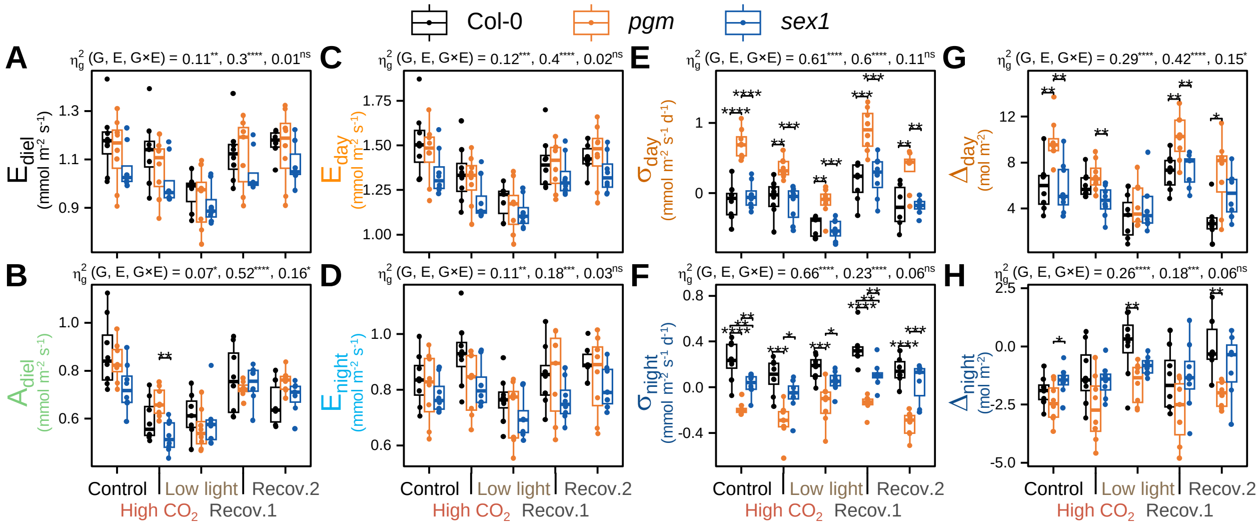


**Supplemental Figure S3.** Comparison of further transpiration parameters for *pgm* and *sex1*. The transpiration dynamics presented in Figure 3 and analysed in Figure 4 were subjected to further analysis: **(A)** mean transpiration over 24-h ($E_{diel}$), **(B)** diel amplitude of transpiration ($A_{diel}$), **(C)** mean transpiration over the daytime ($E_{day}$), **(D)** mean transpiration over the nightime ($E_{night}$), **(E)** average slope of change in transpiration throughout the daytime after 30-min illumination ($\sigma_{day}$), **(F)** average slope of change in transpiration throughout the nighttime after 30-min darkness ($\sigma_{night}$), **(G)** cumulative transpiration dynamics above the daytime trend ($\Delta_{day}$) and **(H)** cumulative transpiration dynamics below the nighttime trend ($\Delta_{night}$). For each parameter, a two-way ANOVA was done and the effect sizes ($\eta^2_g$) of the genotype (G), of the environment (E) and of their interaction (G×E), as well as their respective significance level, are reported above each corresponding plot. Pairwise t-tests were systematically done to assess the differences between genotypes in a given environment, and the differences are reported on the plot when significant. Significance codes for the two-way ANOVA effect sizes and pairwise t-tests are like those in Supplemental Figure S1. Supports Figures 3 and 4.

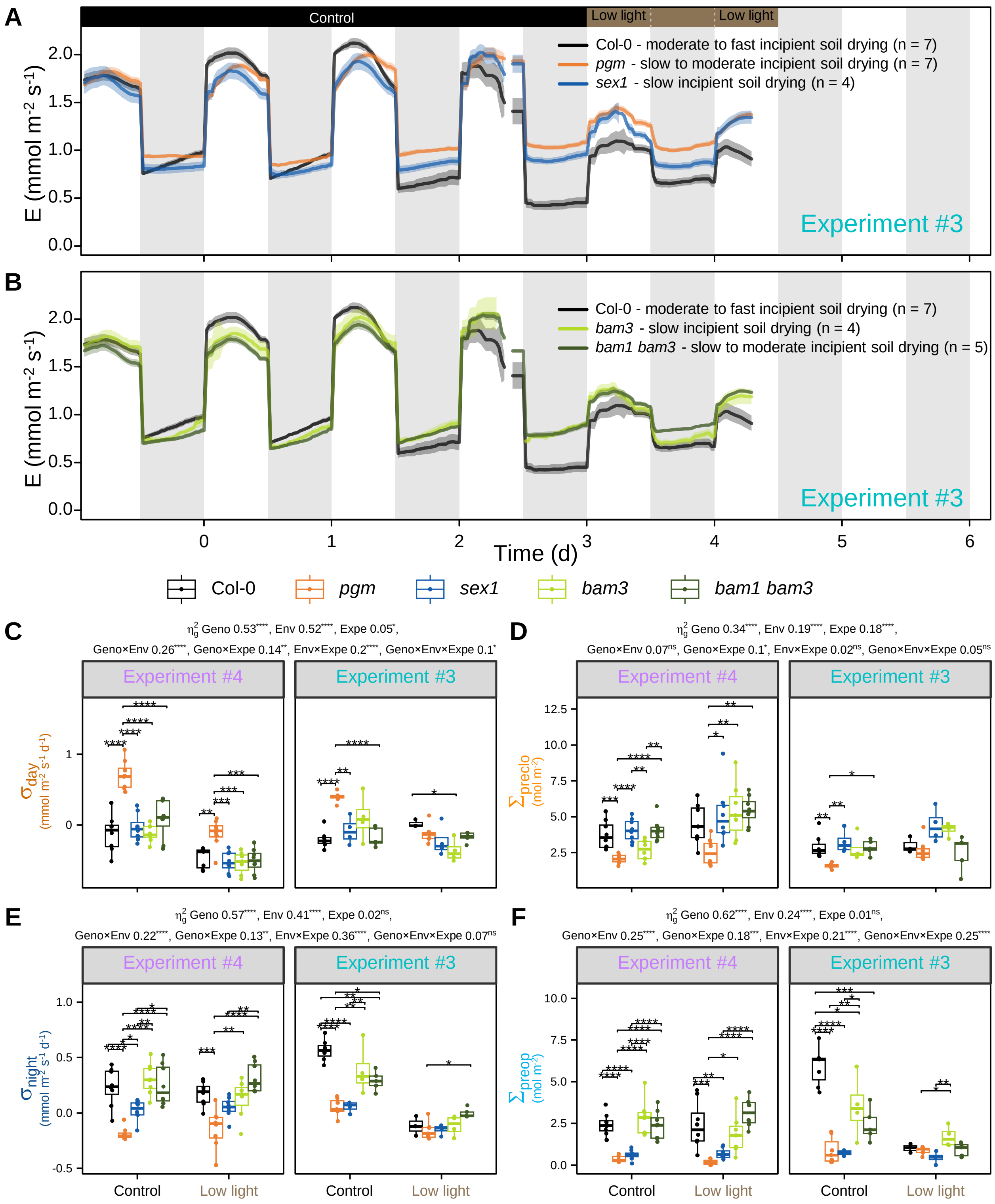


**Supplemental Figure S4.** Analysis of diel transpiration of several starch mutants from the pilot experiment #3 and statistical comparison with the main experiment #4. **(A)** and **(B)** Diel dynamics of transpiration rate of the wild type Col-0 compared to *pgm* and *sex1* **(A)** and to *bam3* and *bam1 bam3* **(B)** in the same conditions as in Figure 2D (experiment #3). The colour-shaded areas around the mean lines represent the means ± SE. **(C)** to **(F)** Boxplots of selected parameters extracted from the measured data: **(C)** $\sigma_{day}$, **(D)** $\Sigma_{preclo}$, **(E)** $\sigma_{night}$ and **(F)** $\Sigma_{preop}$. To avoid excessive differences in soil water content, the control time-period was restricted to the early kinetics, and the wild-type plants categorized as fast soil drying were discarded for the low light period. For each parameter, a three-way ANOVA was done and the effect sizes ($\eta^2_g$) of the genotype (G), of the environment (E), of the experiment (Expe) and of their double and triple interactions, as well as their respective significance level, are reported above each corresponding plot. Pairwise t-tests were systematically done to assess the differences between genotypes in a given environment and experiment, and the differences are reported on the plot when significant. Abbreviations for parameters and significance codes for the three-way ANOVA effect sizes and pairwise t-tests are like those in Supplemental Figure S1. Supports Figures 3, 4 and Supplemental Figure S2.

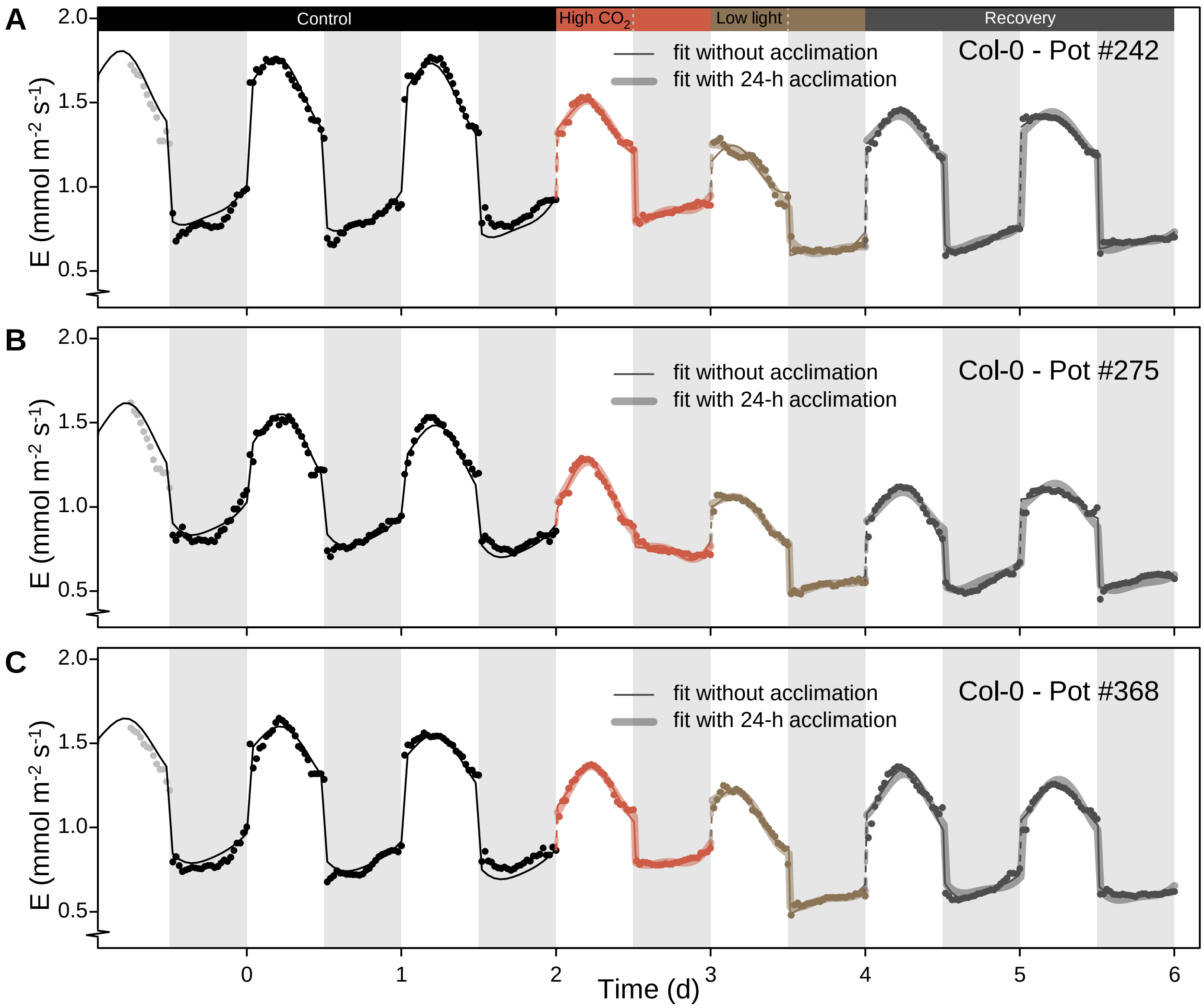


**Supplemental Figure S5.** Examples of individual fits in Col-0. The harmonic model presented in Figure 5 was fitted to the data (points). The initial model was then augmented with a 24-h 'acclimation pulse' when the environment changed, and fitted to the relevant data. Panels **(A)**, **(B)** and **(C)** show the results of the fits for three independent plants of experiment #4 (same conditions as in Figure 3). In each panel, the thin line shows the fitted initial model while the thick, semi-transparent line shows the augmented model. The dashed lines indicate a change in the fit and thus a new set of parameters: data of the previous night did not constrain the fit of the next day when the environment changed. Note that the consecutive recovery days were fitted separately using the initial model, and jointly using the augmented model. The grey points show data of the initial, incomplete daytime period in control conditions that were not used for the fit. Supports Figure 5.

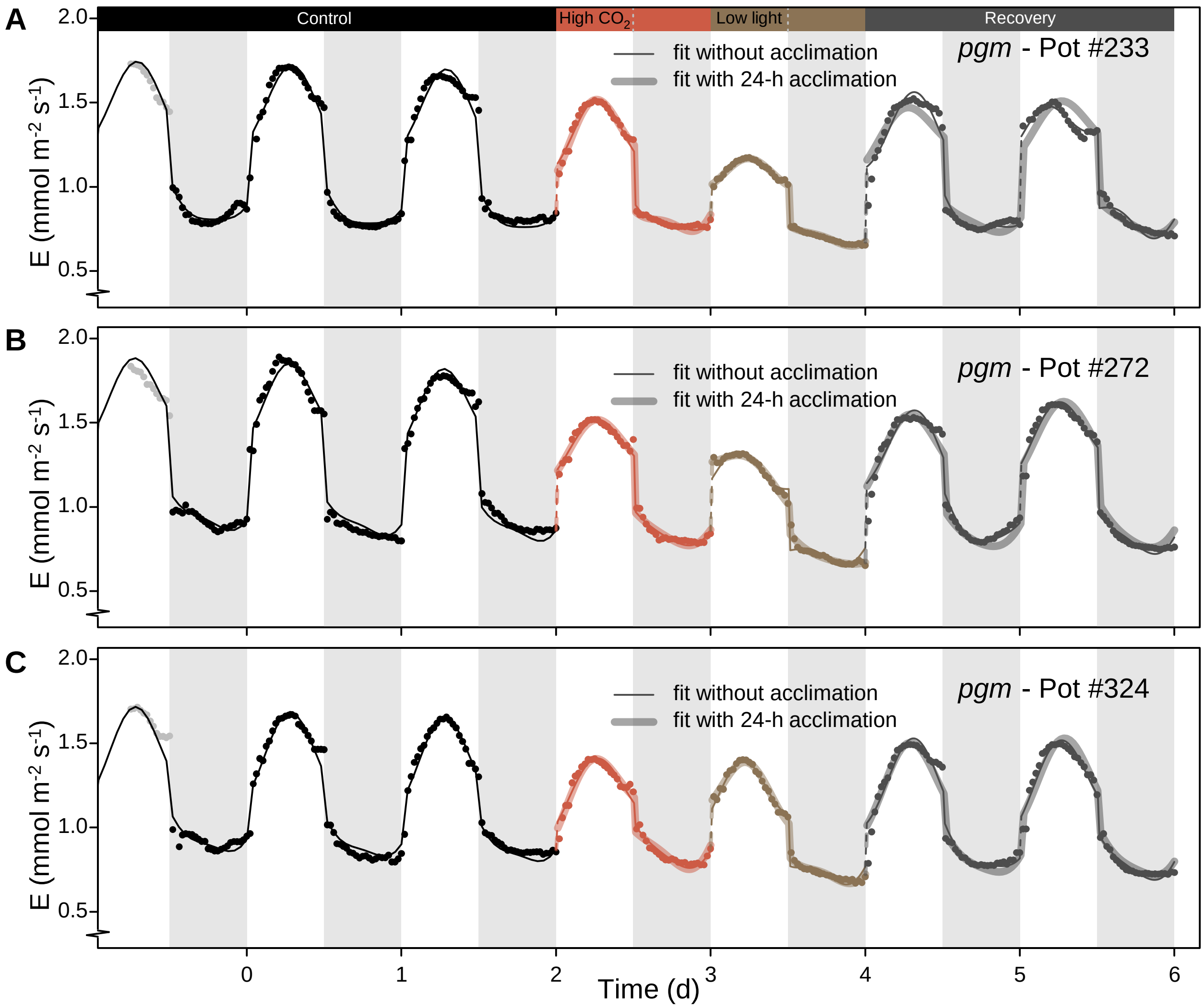


**Supplemental Figure S6.** Examples of individual fits in *pgm*. The harmonic model presented in Figure 5 was fitted to the data (points). The initial model was then augmented with a 24-h 'acclimation pulse' when the environment changed, and fitted to the relevant data. Panels **(A)**, **(B)** and **(C)** show the results of the fits for three independent plants of experiment #4 (same conditions as in Figure 3). In each panel, the thin line shows the fitted initial model while the thick, semi-transparent line shows the augmented model. The dashed lines indicate a change in the fit and thus a new set of parameters: data of the previous night did not constrain the fit of the next day when the environment changed. Note that the consecutive recovery days were fitted separately using the initial model, and jointly using the augmented model. The grey points show data of the initial, incomplete daytime period in control conditions that were not used for the fit. Supports Figure 5.

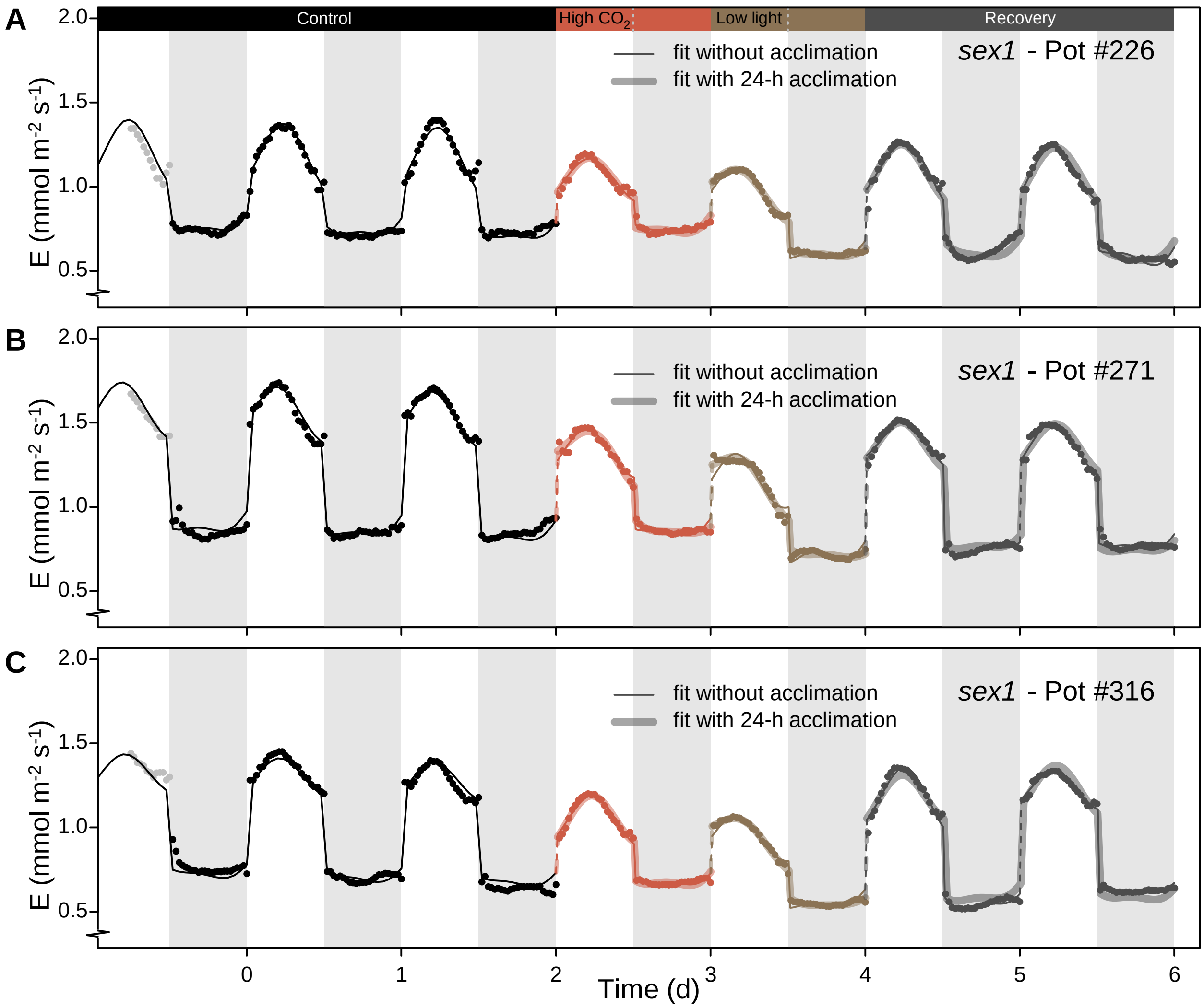


**Supplemental Figure S7.** Examples of individual fits in *sex1*. The harmonic model presented in Figure 5 was fitted to the data (points). The initial model was then augmented with a 24-h 'acclimation pulse' when the environment changed, and fitted to the relevant data. Panels **(A)**, **(B)** and **(C)** show the results of the fits for three independent plants of experiment #4 (same conditions as in Figure 3). In each panel, the thin line shows the fitted initial model while the thick, semi-transparent line shows the augmented model. The dashed lines indicate a change in the fit and thus a new set of parameters: data of the previous night did not constrain the fit of the next day when the environment changed. Note that the consecutive recovery days were fitted separately using the initial model, and jointly using the augmented model. The grey points show data of the initial, incomplete daytime period in control conditions that were not used for the fit. Supports Figure 5.

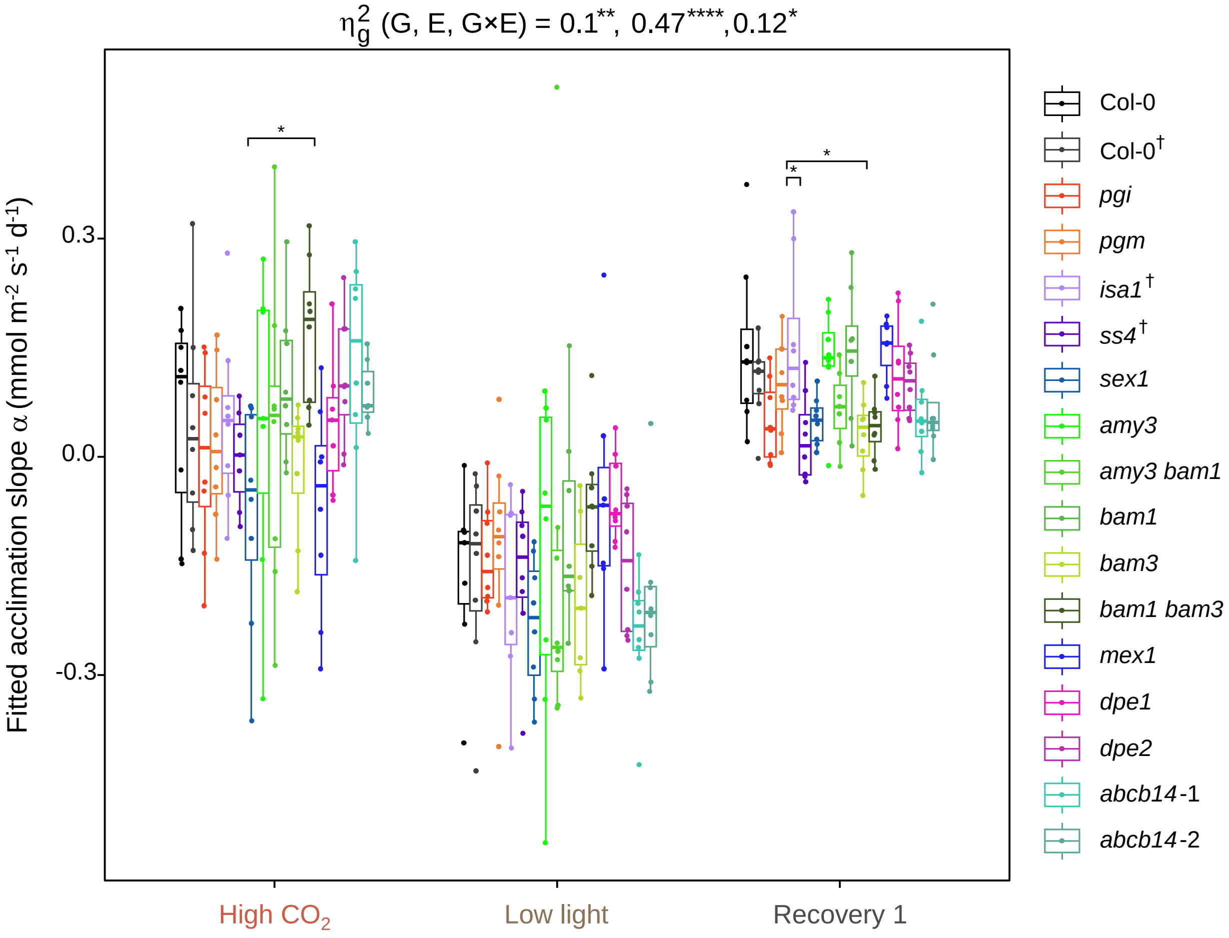


**Supplemental Figure S8.** Effect of the environment and the genotype on the fitted acclimation slope. The acclimation slope parameter a was obtained for high $CO_2$, low light and the first diel cycle of recovery for all genotypes in experiment #4. Significance codes for the two-way ANOVA effect sizes ($\eta^2_g$) and pairwise t-tests are like those in Supplemental Figure S1. Supports Figure 5.

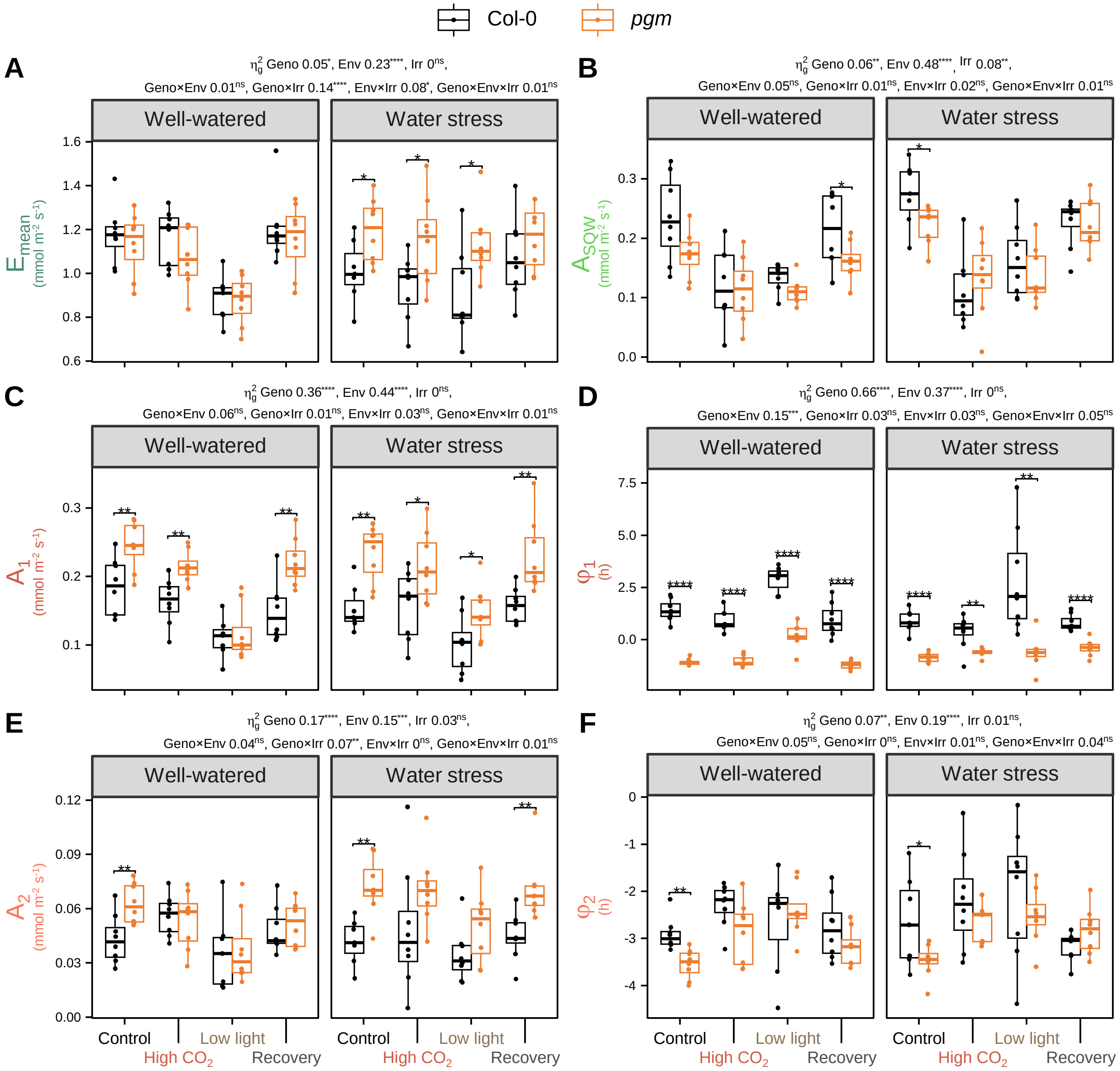


**Supplemental Figure S9.** Effect of the irrigation regime, the environment and the genotype on the fitted parameters. Statistical comparison of the fitted parameters for Col-0 and *pgm* in experiment #4. **(A)** to **(F)** Boxplots of fitted parameters: **(A)** $E_{mean}$, **(B)** $A_{SQW}$, **(C)** $A_1$, **(D)** $\varphi_1$, **(E)** $A_2$ and **(I)** $\varphi_2$. For each parameter, a three-way ANOVA was done and the effect sizes ($\eta^2{}_g$) of the genotype (G), of the environment (E), of the irrigation regime (Irr) and of their double and triple interactions, as well as their respective significance level, are reported above each corresponding plot. Pairwise t-tests were systematically done to assess the differences between genotypes in a given environment and irrigation regime, and the differences are reported on the plot when significant. Abbreviations for parameters and significance codes for the three-way ANOVA effect sizes and pairwise t-tests are like those in Supplemental Figure S1. Supports Figures 2, 5 and Supplemental Figure S2.

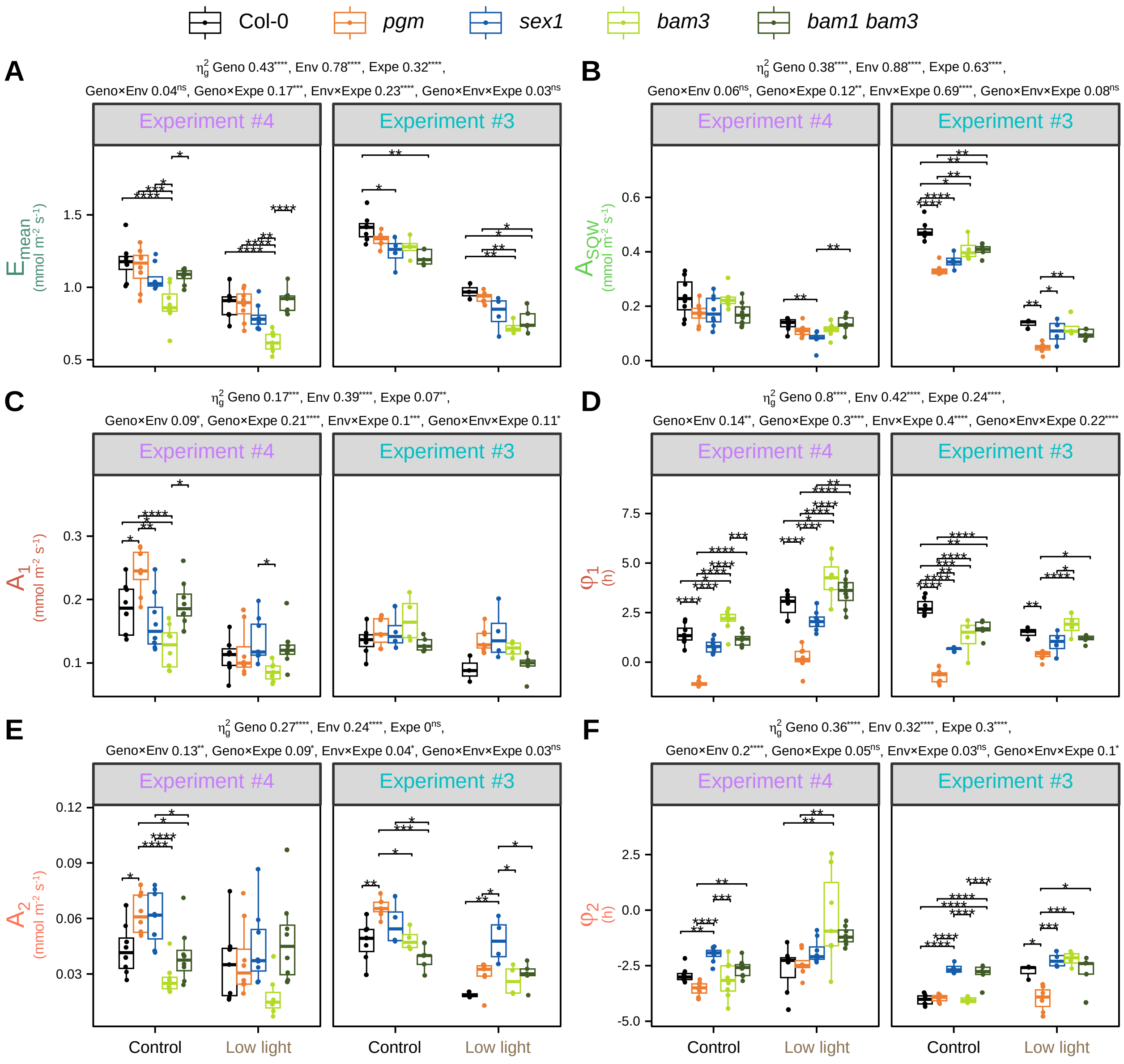


**Supplemental Figure S10.** Effect of the experiment, the environment and the genotype on the fitted parameters. Statistical comparison of the fitted parameters for Col-0 and several starch mutants under control conditions or low light in experiments #3 and #4. **(A)** to **(F)** Boxplots of fitted parameters: **(A)** $E_{mean}$, **(B)** $A_{SQW}$, **(C)** $A_1$, **(D)** $\varphi_1$, **(E)** $A_2$ and **(I)** $\varphi_2$. To avoid excessive differences in soil water content, the control time-period was restricted to the early kinetics, and the wild-type plants categorized as fast soil drying were discarded for the low light period. For each parameter, a three-way ANOVA was done and the effect sizes ($\eta^2_g$) of the genotype (G), of the environment (E), of the experiment (Expe) and of their double and triple interactions, as well as their respective significance level, are reported above each corresponding plot. Pairwise t-tests were systematically done to assess the differences between genotypes in a given environment and experiment, and the differences are reported on the plot when significant. Abbreviations for parameters and significance codes for the three-way ANOVA effect sizes and pairwise t-tests are like those in Supplemental Figure S1. Supports Figures 2, 5 and Supplemental Figures S2 and S4.

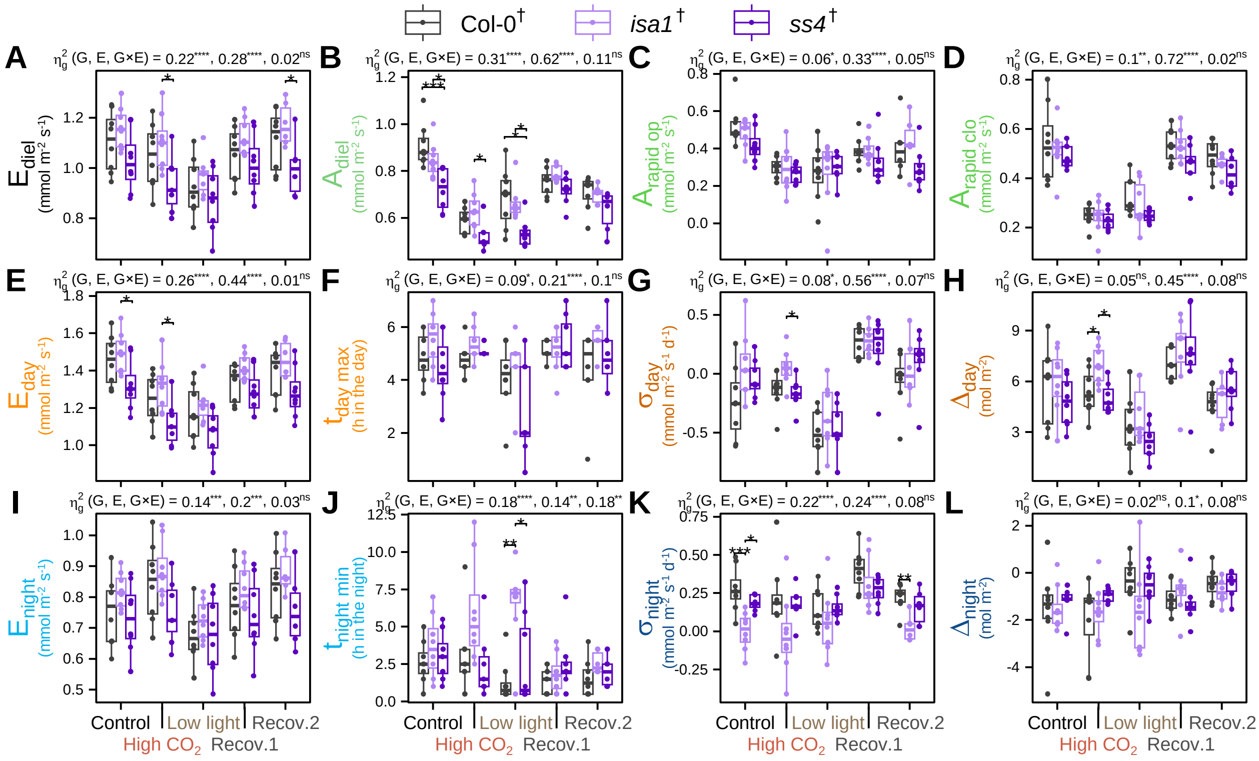

**Supplemental Figure S11.** Comparison of further transpiration parameters for *isa1* and *ss4*. The transpiration dynamics presented in Figure 6 were subjected to further analysis: **(A)** $E_{diel}$, **(B)** $A_{diel}$, **(C)** $A_{rapid\ op}$, **(D)** $A_{rapid\ clo}$, **(E)** $E_{day}$, **(F)** $t_{day\ max}$, **(G)** $\sigma_{day}$, **(H)** $\Delta_{day}$, **(I)** $E_{night}$, **(J)** $t_{night\ min}$, **(K)** $\sigma_{night}$ and **(L)** $\Delta_{night}$. Abbreviations for parameters and significance codes for the two-way ANOVA effect sizes ($\eta^2_g$) and pairwise t-tests are like those in Supplemental Figure S1. The dagger in Col-0†, *ss4*† and *isa1*† indicates that these lines harbour a luciferase reporter that was not used in this study (Methods). Supports Figure 6.

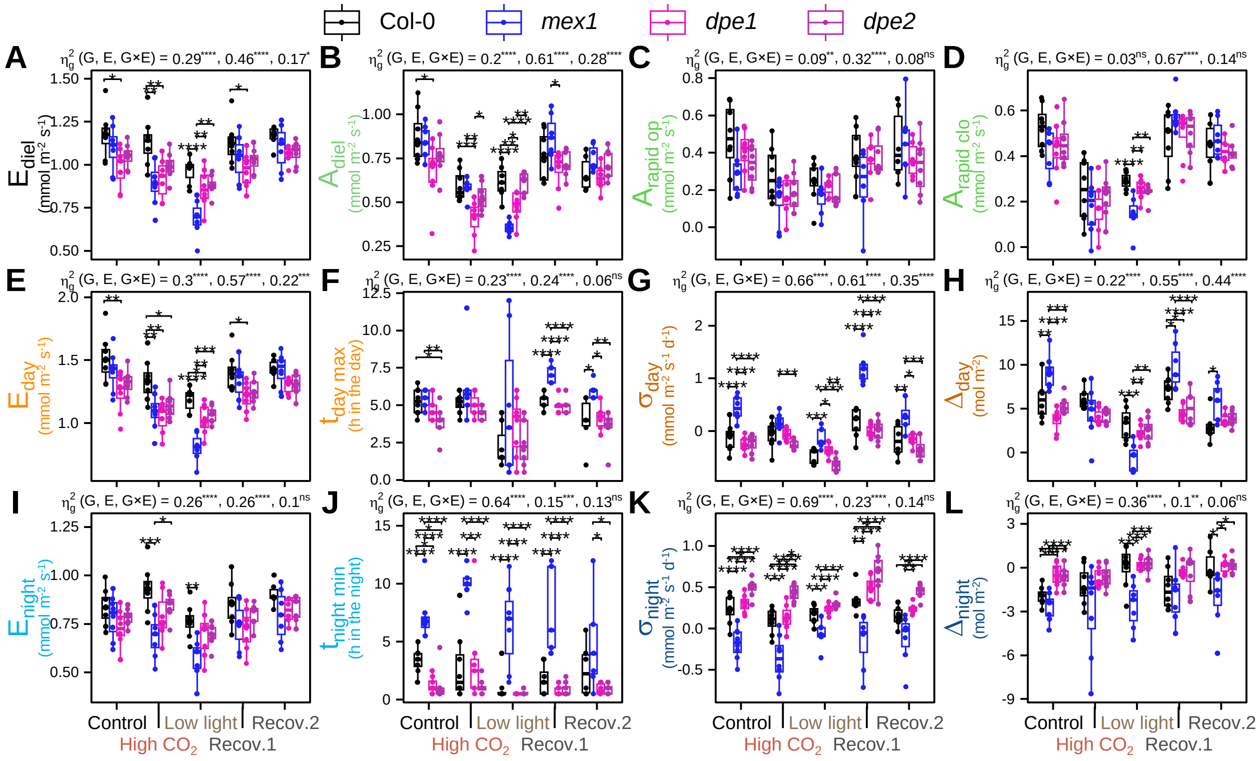


**Supplemental Figure S12.** Comparison of further transpiration parameters for *mex1*, *dpe1* and *dpe2*. The transpiration dynamics presented in Figure 7 were subjected to further analysis: **(A)** $E_{diel}$, **(B)** $A_{diel}$, **(C)** $A_{rapid\ op}$, **(D)** $A_{rapid\ clo}$, **(E)** $E_{day}$, **(F)** $t_{day\ max}$, **(G)** $\sigma_{day}$, **(H)** $\Delta_{day}$, **(I)** $E_{night}$, **(J)** $t_{night\ min}$, **(K)** $\sigma_{night}$ and **(L)** $\Delta_{night}$. Abbreviations for parameters and significance codes for the two-way ANOVA effect sizes ($\eta^2_g$) and pairwise t-tests are like those in Supplemental Figure S1. Supports Figure 7.

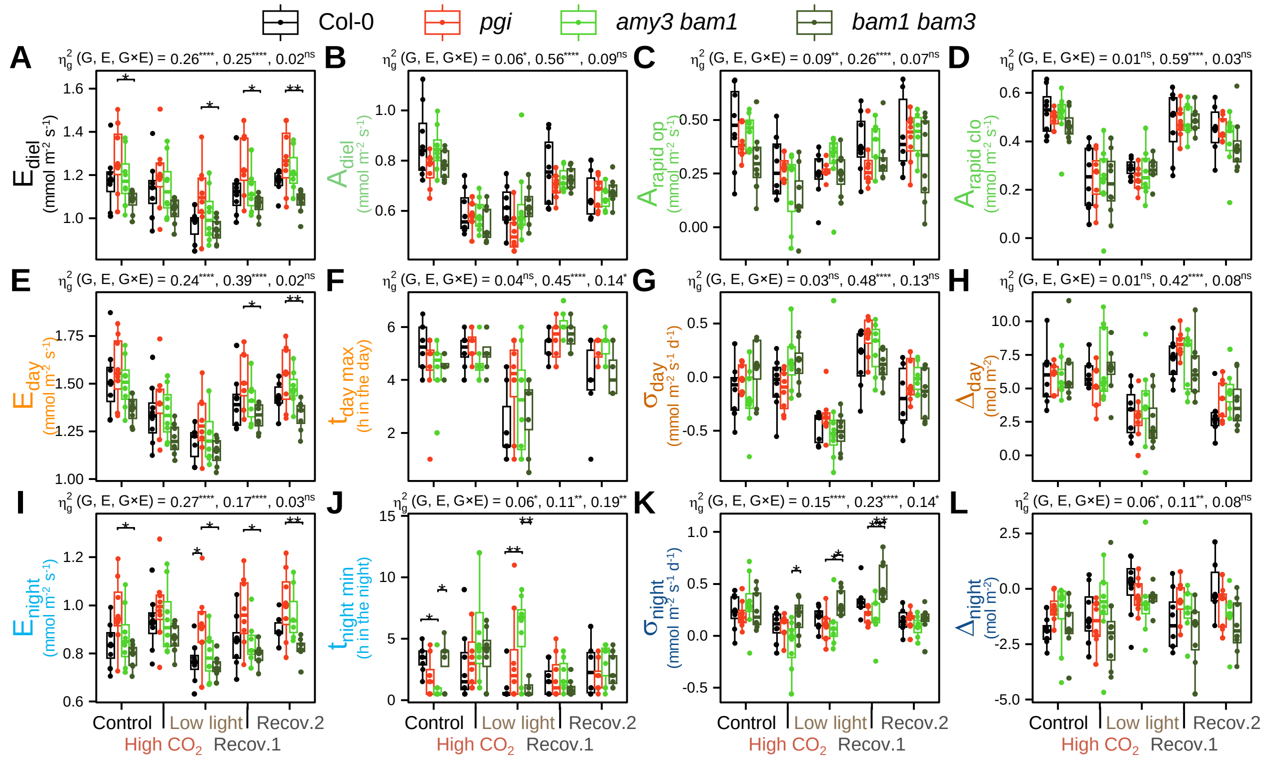


**Supplemental Figure S13.** Comparison of further transpiration parameters for *pgi*, *amy3 bam1* and *bam1 bam3*. The transpiration dynamics presented in Figure 8 were subjected to further analysis: **(A)** $E_{diel}$, **(B)** $A_{diel}$, **(C)** $A_{rapid\ op}$, **(D)** $A_{rapid\ clo}$, **(E)** $E_{day}$, **(F)** $t_{day\ max}$, **(G)** $\sigma_{day}$, **(H)** $\Delta_{day}$, **(I)** $E_{night}$, **(J)** $t_{night\ min}$, **(K)** $\sigma_{night}$ and **(L)** $\Delta_{night}$. Abbreviations for parameters and significance codes for the two-way ANOVA effect sizes ($\eta^2_g$) and pairwise t-tests are like those in Supplemental Figure S1. Supports Figure 8.

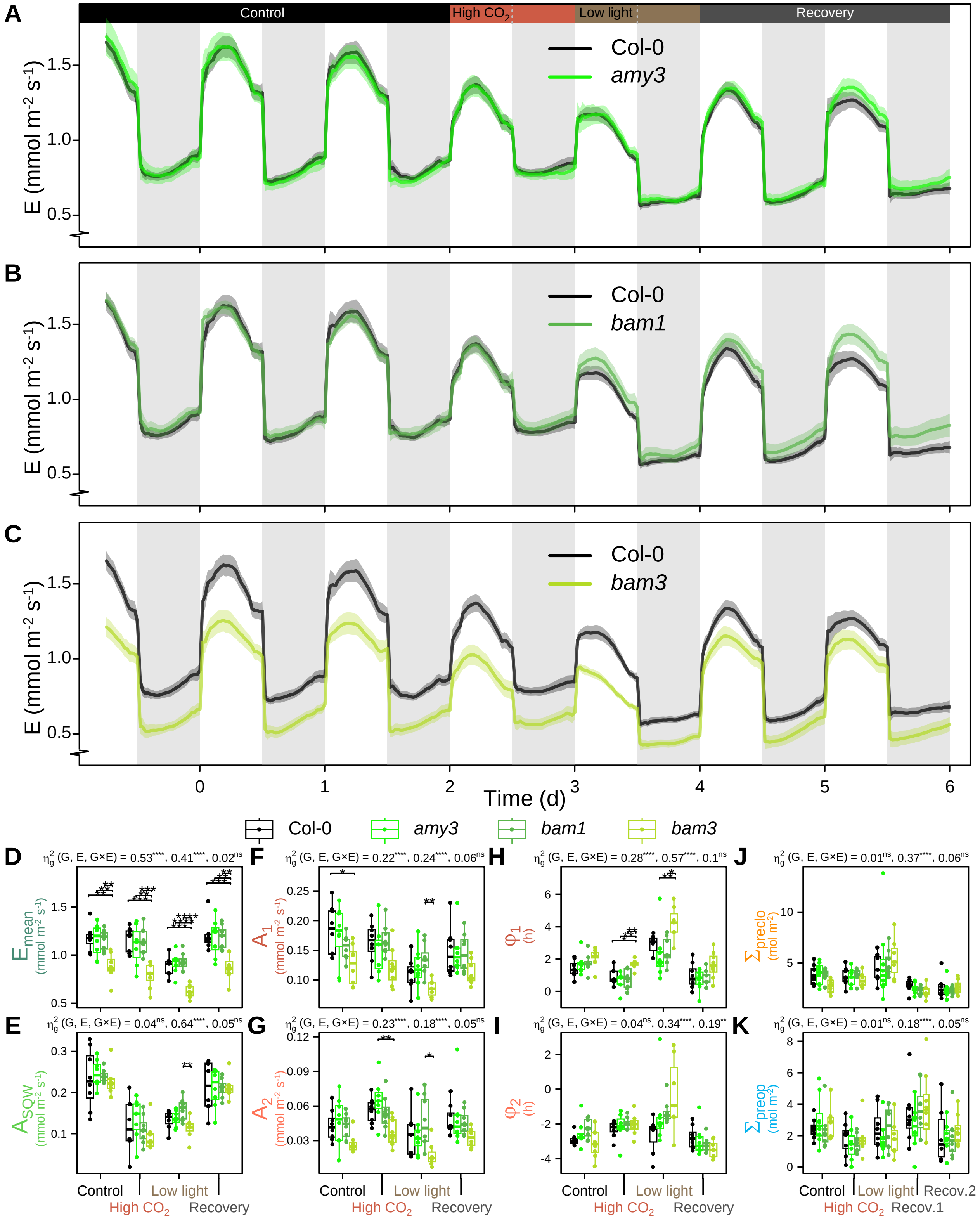


**Supplemental Figure S14.** Analysis of diel transpiration of *amy3*, *bam1* and *bam3* single mutants. **(A)** to **(C)** Diel dynamics of transpiration rate of the wild type Col-0 compared to *amy3* **(A)**, *bam1* **(B)** and *bam3* **(C)** in the same conditions as in Figure 3. The colour-shaded areas around the mean lines represent the means ± SE. **(D)** to **(I)** Boxplots of fitted parameters: **(D)** $E_{mean}$, **(E)** $A_{SQW}$, **(F)** $A_1$, **(G)** $A_2$, **(H)** $\varphi_1$ and **(I)** $\varphi_2$. **(J)** and **(K)** Boxplots of selected parameters extracted from the measured data: **(J)** $S_{preclo}$ and **(K)** $S_{preop}$. Abbreviations for parameters and significance codes for the two-way ANOVA effect sizes ($\eta^2_g$) and pairwise t-tests are like those in Supplemental Figure S1. Supports Figure 8.

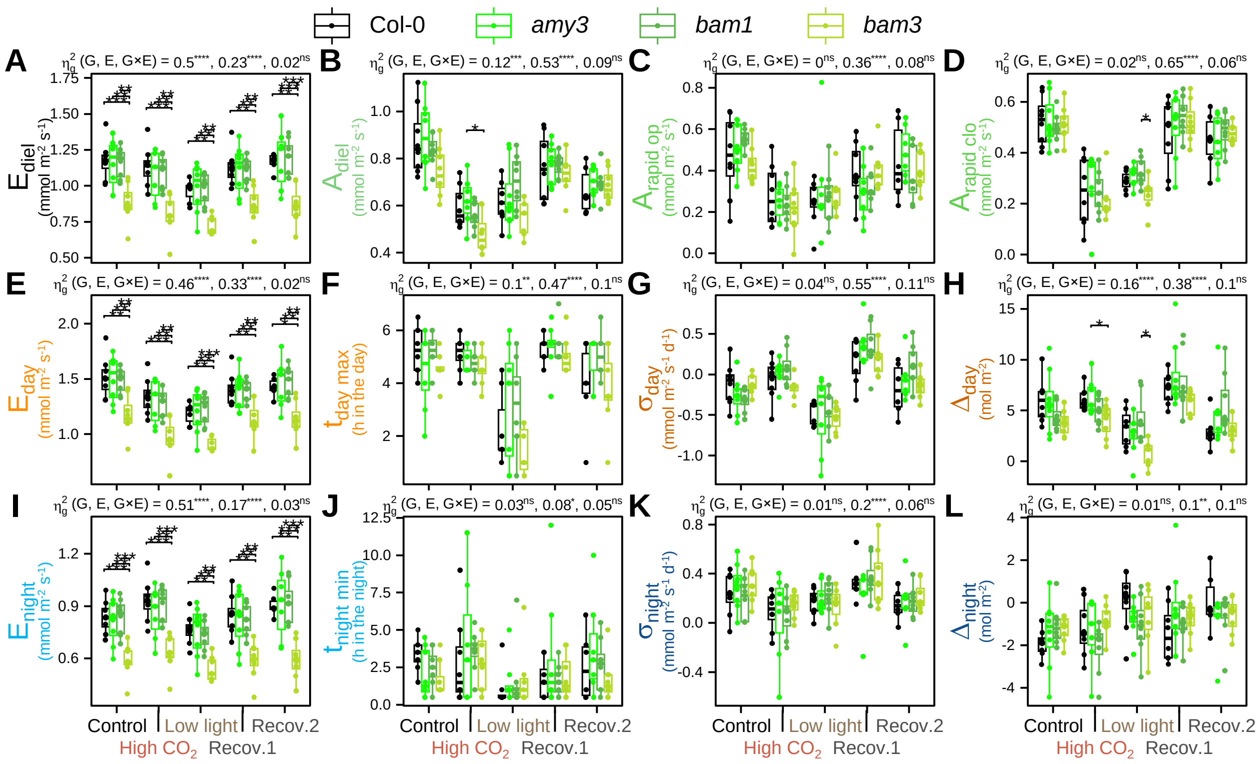

**Supplemental Figure S15.** Comparison of further transpiration parameters for *amy3*, *bam1* and *bam3*. The transpiration dynamics presented in Supplemental Figure S14 were subjected to further analysis: **(A)** $E_{diel}$, **(B)** $A_{diel}$, **(C)** $A_{rapid\ op}$, **(D)** $A_{rapid\ clo}$, **I** $E_{day}$, **(F)** $t_{day\ max}$, **(G)** $\sigma_{day}$, **(H)** $\Delta_{day}$, **(I)** $E_{night}$, **(J)** $t_{night\ min}$, **(K)** $\sigma_{night}$ and **(L)** $\Delta_{night}$. Abbreviations for parameters and significance codes for the two-way ANOVA effect sizes ($\eta^2_g$) and pairwise t-tests are like those in Supplemental Figure S1. Supports Figure 8 and Supplemental Figure S14.

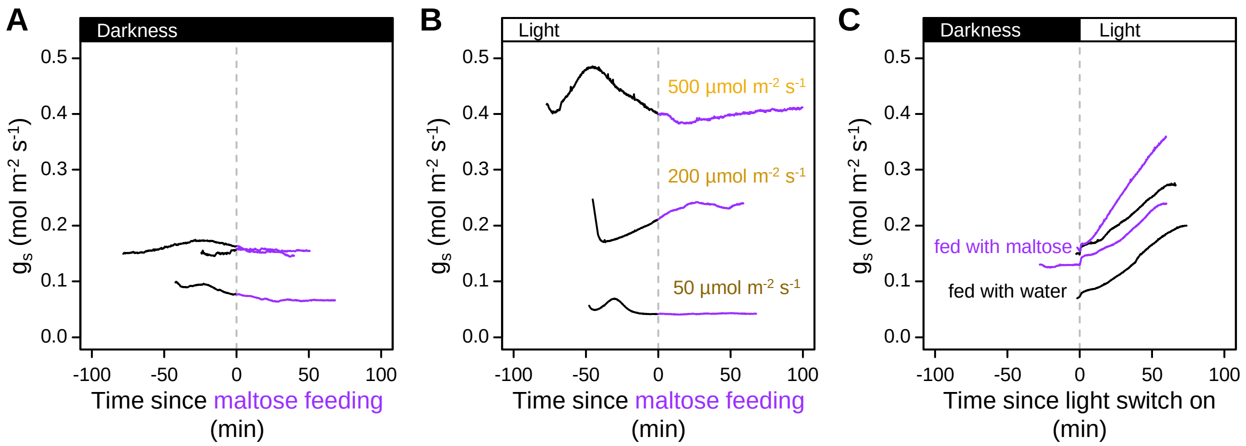


**Supplemental Figure S16.** Effect of exogenous maltose on stomatal conductance of Col-0. Gas-exchange experiments were performed on detached leaves of Col-0 that were xylem-fed with a high concentration of maltose (1 mM). The dynamics of stomatal conductance ($g_s$) was monitored using a Li-Cor 6400. **(A)** Maltose-feeding in darkness at the end of the night. **(B)** Maltose-feeding in the light (50, 200 or 500 µmol $m^{-2}$ $s^{-1}$) during the daytime. **(C)** Leaves fed with water (control) or maltose at the end of the night, subjected to extended darkness, enclosed in the Li-Cor 6400 chamber and then exposed to light (500 µmol $m^{-2}$ $s^{-1}$). Supports Figures 7 and 8.

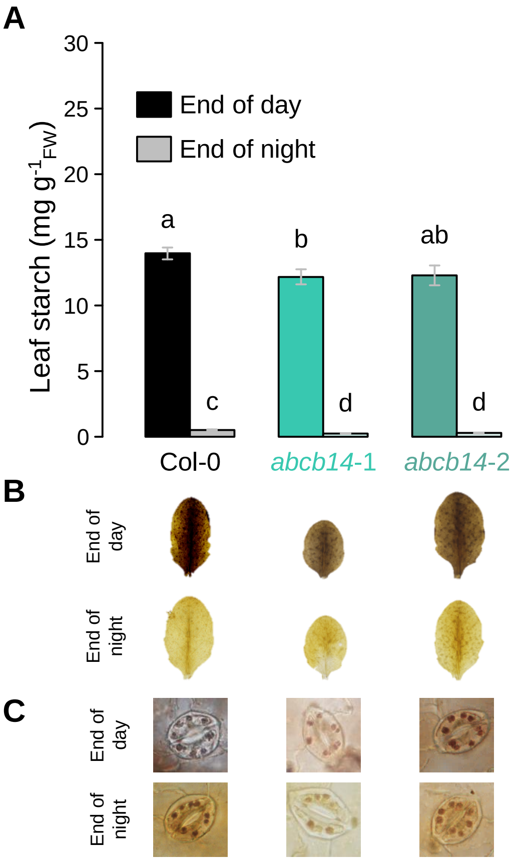


**Supplemental Figure S17.** Characterization of the starch patterns in *abcb14*-1 and *abcb14*-2. **(A)** Leaf starch content determined at the end of the day and night periods in control conditions. Error bars are means ± SE. Letters denote significant differences after a Kruskal-Wallis test (α = 5%) followed by multiple comparisons of ranks. **(B)** and **(C)** Iodine staining of starch on entire leaves **(B)** or individual guard cells **(C)** at the end of the day (top rows) or night (bottom rows). The data and images for Col-0 are the same as the ones used in Figure 1. Supports Figure 9 and Supplemental Figure S18.

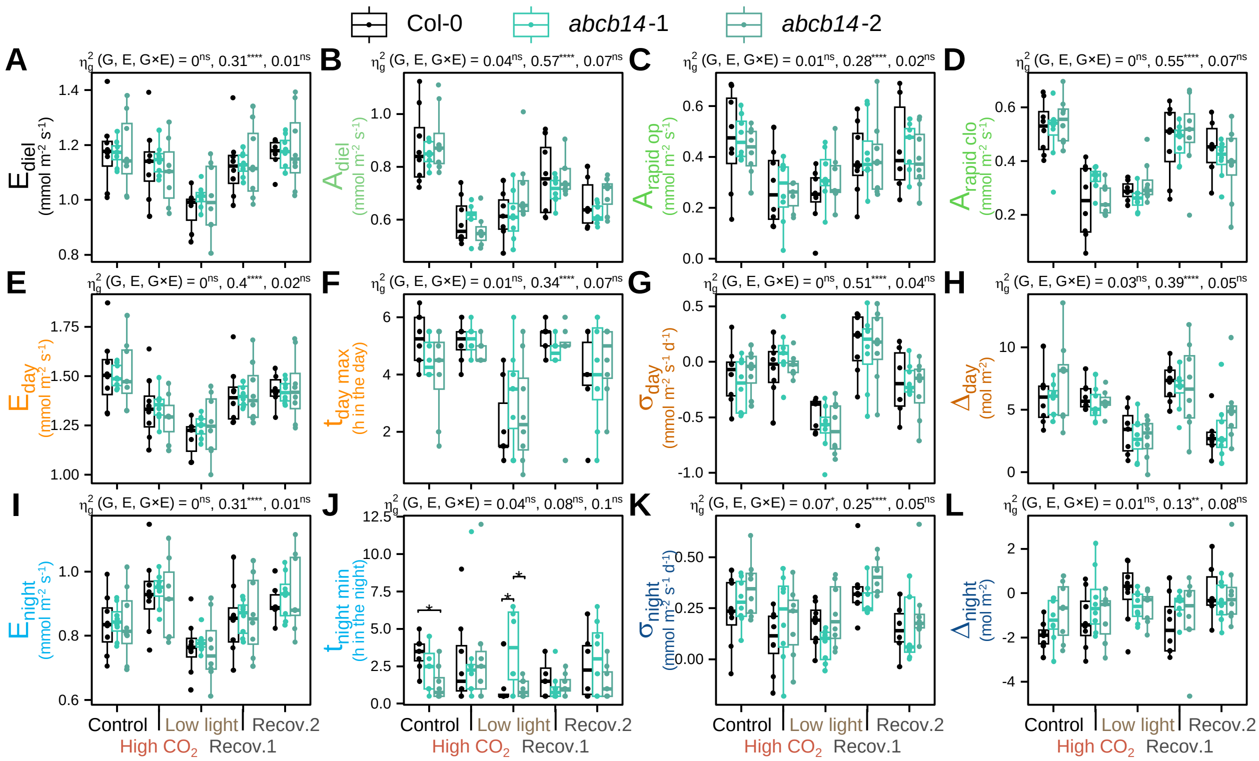


**Supplemental Figure S18.** Comparison of further transpiration parameters for *abcb14*-1 and *abcb14*-2. The transpiration dynamics presented in Figure 9 were subjected to further analysis: **(A)** $E_{diel}$, **(B)** $A_{diel}$, **(C)** $A_{rapid\ op}$, **(D)** $A_{rapid\ clo}$, **(E)** $E_{day}$, **(F)** $t_{day\ max}$, **(G)** $\sigma_{day}$, **(H)** $\Delta_{day}$, **(I)** $E_{night}$, **(J)** $t_{night\ min}$, **(K)** $\sigma_{night}$ and **(L)** $\Delta_{night}$. Abbreviations for parameters and significance codes for the two-way ANOVA effect sizes ($\eta^2_g$) and pairwise t-tests are like those in Supplemental Figure S1. Supports Figure 9.
